# Maintaining transcriptome solubility constrains mRNA sequence composition

**DOI:** 10.1101/2025.09.19.677370

**Authors:** Marco Todisco, Christalyn Ausler, Ankur Jain

**Author notes:** **Author Contributions:** M.T. implemented the simulation code, performed experiments and data analysis. C.A. performed mRNA extraction. All authors contributed to project conception and manuscript writing. **Competing Interest Statement:** The authors declare that they have no competing financial interests.

## Abstract

RNA is built from a four-nucleotide alphabet. Complementary sequences inevitably arise, creating pervasive opportunities for promiscuous RNA–RNA interactions. Here, we show that this chemistry makes the transcriptome intrinsically prone to self-association. We simulated the simultaneous interactions of ~7,500 mRNAs representing the *E. coli* transcriptome at physiological concentrations. These large-scale simulations predict widespread, dynamic clustering driven by RNA alone and organized by long, multivalent transcripts. Purified mRNA recapitulates this behavior *in vitro*, with aggregate composition mirroring model predictions. Strikingly, native mRNA sequences are markedly less prone to self-association than matched randomized controls: they fold more stably, expose shorter single-stranded regions, and form weaker intermolecular contacts. Similar signatures are observed in abundant human mRNAs, suggesting that evolution has shaped coding sequences to minimize self-association. These findings identify transcriptome solubility as an unrecognized constraint on mRNA sequence evolution and provide a framework for understanding how cells keep their transcriptomes dispersed and functional.

## Introduction

The base-pairing chemistry that underlies RNA folding and regulation also creates opportunities for unintended RNA-RNA interactions. Because RNA is built from a four-letter alphabet, short complementary sequence tracts arise frequently by chance (1), making large RNA pools intrinsically prone to self-association (2). Indeed, RNA’s tendency to form promiscuous intermolecular base pairs has been appreciated for decades: standard protocols for RNA analysis routinely employ strong denaturants to prevent aggregation (3–6). Whether this tendency poses a meaningful constraint on cellular transcriptomes remains unknown.

This question is especially relevant for messenger RNAs. Although mRNAs are often viewed primarily as carriers of coding information, in the cytoplasm they exist as a concentrated and compositionally diverse pool of molecules that are capable of intramolecular folding and intermolecular pairing. Recent transcriptome-wide studies have revealed extensive RNA-RNA associations *in vivo* (7, 8). RNA-RNA interactions have also been implicated in the formation of cellular condensates such as stress granules (9, 10), TIS granules (11), germ granules (12), and disease-associated RNA aggregates (13–15). Notably, purified yeast RNA, freed from proteins, readily self-assembles into structures whose RNA composition resembles that of cellular stress granules (9). Taken together, these observations suggest that RNA-RNA interactions *in vivo* are both plausible and potentially consequential. A natural question is whether the self-association observed *in vitro* also occurs inside cells. Under normal growth conditions, most mRNAs remain dispersed, suggesting that cellular factors prevent aggregation (16–18). But how much of this dispersal reflects the intrinsic properties of mRNA sequences themselves, and how much requires active suppression by ribosomes, RNA-binding proteins, and helicases? Answering this question requires first defining a baseline: how much aggregation would RNA chemistry alone produce at physiological concentrations?

RNA folding and pairwise hybridization have been studied extensively for decades (19, 20), establishing a detailed understanding of secondary structure and duplex formation *in vitro*. However, these studies typically consider one or a few molecules at a time. Extending these principles to predict the collective behavior of thousands of RNA molecules coexisting at high concentration, as they do in cells, remains challenging (21–28). A quantitative RNA-only baseline for transcriptome-wide mRNA self-association is therefore still lacking.

Here, we establish this baseline. We develop a computational framework that combines RNA folding predictions, pairwise complementarity analysis, and stochastic kinetic simulations to estimate transcriptome-wide intermolecular base pairing when each mRNA is present at its physiological concentration. Applied to *E. coli*, the model predicts that RNA chemistry alone is sufficient to drive widespread, dynamic clustering, organized around long, multivalent mRNAs. Importantly, purified *E. coli* mRNA readily aggregated *in vitro*, and the composition of aggregated mRNA closely matches that predicted by the simulations. This framework also allowed us to ask whether aggregation pressure has shaped sequence evolution. Native mRNA sequences exhibit more stable intramolecular folding, shorter unstructured stretches, and significantly weaker intermolecular interactions than shuffled controls. Similar signatures are also observed in mammalian mRNAs. Together, these findings identify transcriptome solubility as an underappreciated constraint on mRNA sequence evolution and define an RNA-only reference point for understanding how cells keep their mRNA pools dispersed and functional.

## Results

### 1. The transcriptome encodes extensive potential for intermolecular pairing

To determine whether the cellular mRNA pool is intrinsically poised for intermolecular RNA-RNA interactions, we developed a computational framework and applied it to *E. coli*, where both transcript sequences and abundances are well characterized. In living cells, ribosomes, RNA-binding proteins, and helicases modulate RNA base pairing (29–31). To isolate the contribution of RNA chemistry alone, we considered a minimal RNA-only system that defines an upper bound on the sequence-encoded potential for intermolecular pairing. In this setting, intramolecular folding is expected to dominate: each transcript will fold on itself first, and the regions that remain unpaired define what is available for intermolecular contact. The capacity of a transcriptome to self-associate therefore depends on the balance between folded and accessible regions within each mRNA. We set out to map these accessible elements across the entire *E. coli* transcriptome (Fig. 1).

**Figure 1.**
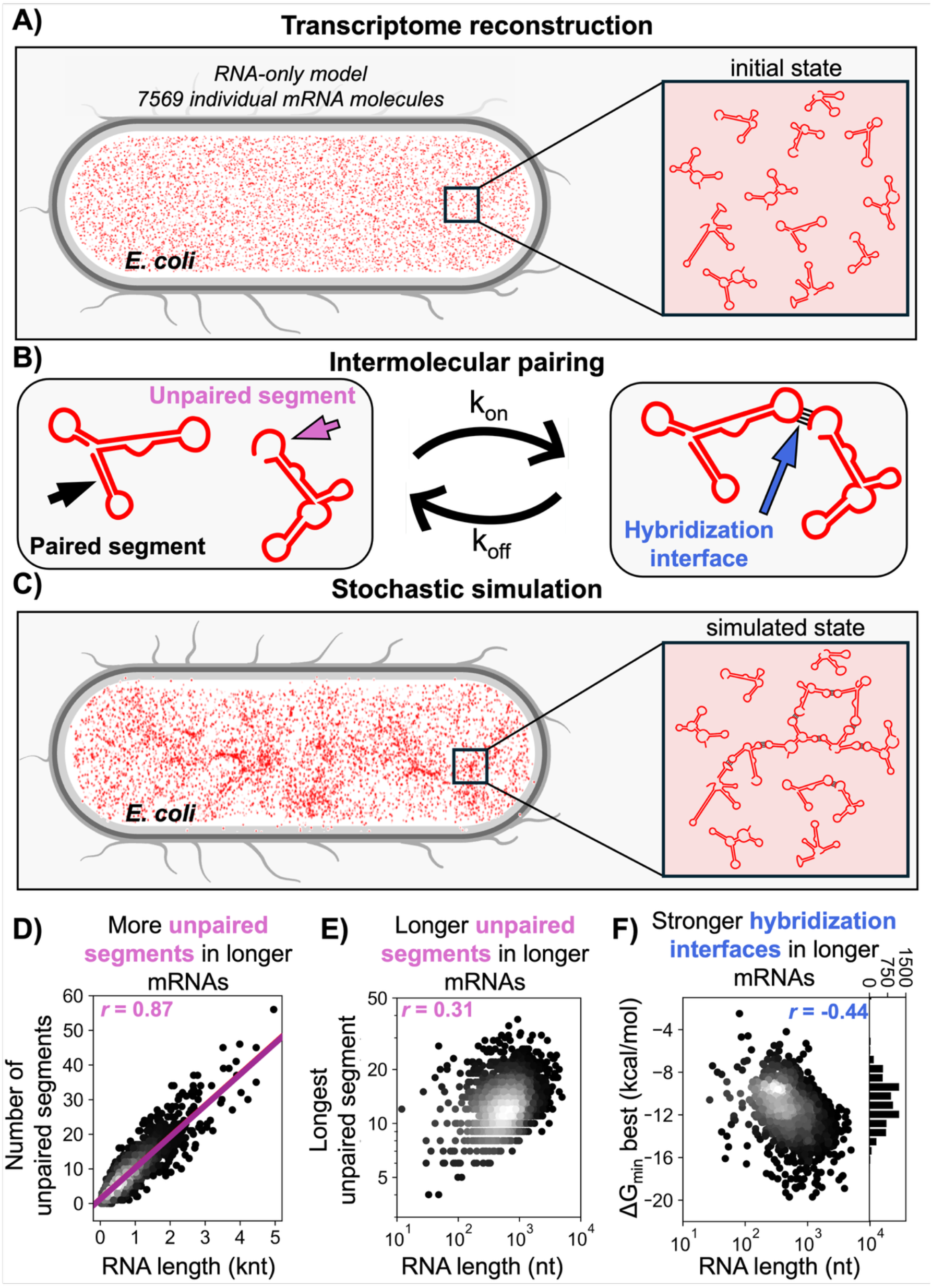
RNA-only framework for estimating transcriptome-wide mRNA self-association. (A) The *E. coli* transcriptome was reconstructed from transcript abundance data and each mRNA was folded *in silico*. (B) Unpaired segments remaining after intramolecular folding were treated as candidate sites for intermolecular hybridization. Schematics illustrate intramolecularly paired regions, unpaired segments available for intermolecular contact, and the hybridization interface between two transcripts. (C) These interaction sites, together with transcript abundances and cellular volume were used to parameterize stochastic simulations of mRNA association and dissociation. (D) Number of unpaired segments (≥7 nt) per transcript as a function of transcript length. Purple line, linear fit; Pearson’s r = 0.87. (E) Length of the longest unpaired segment in each transcript as a function of transcript length; Pearson’s r = 0.31. (F) Most favorable predicted intermolecular hybridization energy (ΔG_min, best_) for each transcript as a function of transcript length; Pearson’s r = −0.44. For each transcript, ΔG_min, best_ denotes the lowest hybridization free energy among all candidate intermolecular interactions involving that transcript. Right, distribution of ΔG_min, best_ values.

We instantiated this framework using *E. coli* grown in standard rich-growth conditions for which both total mRNA content and transcript abundances have been previously estimated (32, 33). Under these conditions, *E. coli* contains an estimated 7,800 mRNA molecules per cell (32) within an intracellular volume of 6.7 × 10^−10^ µL (33), corresponding to a total mRNA concentration of roughly 20 μM. Using published RNA-seq datasets (34), we assigned transcript-specific copy numbers to reconstruct a copy-number-weighted representation of the steady-state mRNA pool. After rounding, the model comprised of 7,569 mRNA molecules representing 4,510 unique coding transcripts (see Supp. Spreadsheet 1). To identify regions available for intermolecular pairing, we folded each transcript independently *in silico* using ViennaRNA (35) and extracted nucleotides with low pairing probability (see Methods A; Fig. S1). Across the transcriptome, approximately 36% of mRNA nucleotides fell below our pairing threshold (probability of base-pairing less than 50%, see Supp. Spreadsheet 2), indicating that a substantial fraction of the transcriptome remains unpaired after intramolecular folding and is therefore potentially accessible for intermolecular interactions.

From these weakly paired nucleotides, we extracted contiguous single-stranded segments of at least 7 nt as candidate interaction sites (Methods B; Fig. S1). We reasoned that shorter segments would be unlikely to form consequential intermolecular duplexes and would greatly expand the search space (Methods B). This analysis identified ~46,000 candidate interaction sites across the 4,510 unique transcripts and ~55,000 sites in the copy-number-weighted cellular pool. These unstructured segments are common, with a median length of 8 nt and occur at a density of approximately 10 per 1,000 nucleotides. Both their number and their maximum length increased with transcript length (Fig. 1D, E), indicating that longer mRNAs are intrinsically more multivalent and expose more, and longer, regions available for intermolecular pairing.

We next asked how readily these accessible regions could find complementary partners elsewhere in the transcriptome. We performed exhaustive pairwise comparisons across all candidate interaction sites and, for each pair of transcripts, identified the most favorable intermolecular interface together with its hybridization energy, *ΔG*_*min*_ (Methods C). This yielded a transcriptome-wide matrix of ~29 million possible pairwise interactions (see Supp. Spreadsheet 3).

The strongest predicted interaction for each transcript became progressively more favorable with transcript length (Fig. 1F), consistent with longer mRNAs harboring more and longer accessible segments. For each transcript, we identified its single most favorable interaction across all possible partners. The median value across transcripts was approximately −11 kcal/mol, comparable to the stability of an 8-bp duplex and similar in magnitude to miRNA seed-region binding (36), while the most favorable interaction approached −20 kcal/mol, comparable to the binding energy of therapeutic antisense oligonucleotides (37).

Favorable interactions were pervasive. Of the ~29 million pairwise comparisons, 28 million yielded negative *ΔG*_*min*_ (see Supp. Spreadsheet 3). Even when considering only strong interactions at a stringent threshold (*ΔG*_*min*_ ≤ –7 kcal/mol or ~12 kT), approximately 2 million interactions remained, involving roughly 38,000 unique sites across the transcriptome and corresponding to ~5 strong-binding sites per mRNA (see Supporting Data). Thus, the native mRNA pool contains extensive latent potential for self-association, with most mRNAs capable of pairing with multiple partners.

As a first approximation, we applied a simple equilibrium self-association model (Supplementary Note 1). This model predicts that roughly 70% of molecules would be incorporated into multimeric complexes at equilibrium. However, this estimate does not account for the fact that the ~2 million strong RNA-RNA interactions identified above are distributed across only ~38,000 unique accessible sites, meaning that most potential interactions are mutually exclusive. Each site instead competes among many alternative partners, with a median of approximately 70 possible binders. Static analysis therefore cannot determine how many contacts can coexist at steady state or whether they are sufficient to generate higher-order assemblies. To address these questions, we turned to stochastic kinetic simulations that explicitly track binding and unbinding events across the transcriptome.

### 2. Stochastic simulations predict widespread, dynamic clustering of the transcriptome

To determine whether the extensive interaction potential identified above is sufficient to drive collective self-association, we performed stochastic kinetic simulations of transcriptome-wide RNA-RNA binding and unbinding. We used a Gillespie algorithm-based framework (38) where each event corresponds either to hybridization between two accessible sites or to dissociation of an existing duplex (Methods D). Hybridization rates (*k*_*on*_) were fixed at *k*_*on*_ = 10^7^ M^−1^s^−1^, consistent with nucleation-limited RNA hybridization (27). Dissociation rates (*k*_*off*_) for each pair were computed from *k*_*on*_ and *ΔG*_*min*_ using detailed balance. Notably, varying *k*_*on*_ over a range of values rescaled the dynamics proportionally without altering steady-state properties (Supplementary Note 2, Fig. S2). To keep the simulations computationally tractable, we included only strong interactions (*ΔG*_*min*_ ≤ −7 kcal/mol). This makes our estimates conservative, because weaker interactions were omitted (Supplementary Note 3, Fig. S3).

Our simulations revealed that the transcriptome is intrinsically poised to form a dynamic network of intermolecular contacts. At steady state, the mRNA pool contained approximately 2,500 concurrent intermolecular interactions (Fig. 2A). About 40% of mRNA molecules engaged in at least one intermolecular contact at any given time, with frequent hybridization and dehybridization events. Because each transcript can have multiple unstructured segments, they can bind several partners simultaneously. This intrinsic multivalency drove the spontaneous formation of higher-order RNA assemblies (Fig. 2B, C).

**Figure 2.**
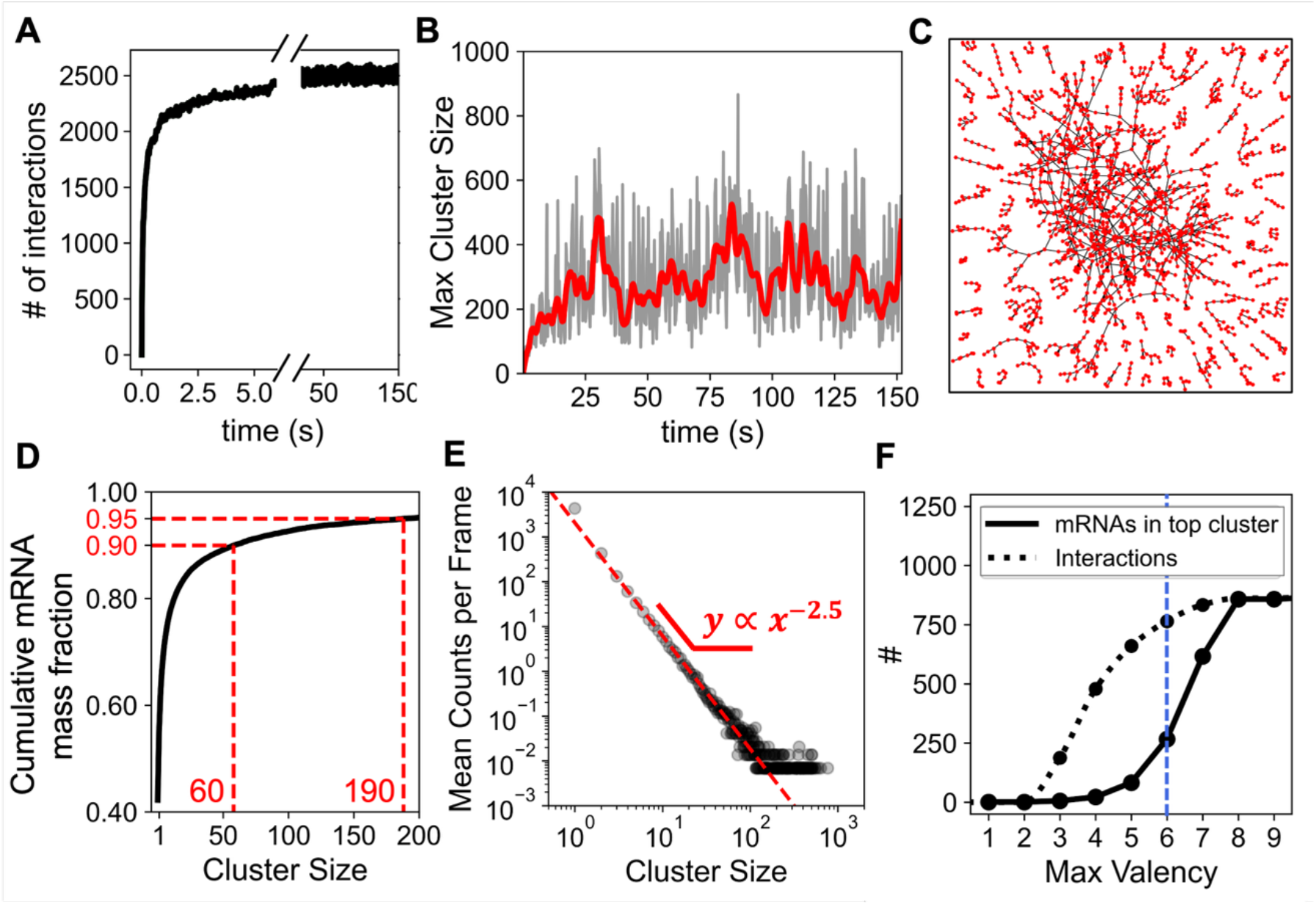
Simulations predict dynamic transcriptome-wide mRNA assemblies. (A) Total number of realized intermolecular mRNA–mRNA interactions over time in a representative simulation, showing rapid establishment of a steady state with ~2,500 concurrent interactions. (B) Size of the largest cluster over time. Large assemblies form and dissolve dynamically, with the largest cluster transiently reaching ~900 mRNAs. Gray, instantaneous values; red, moving average. (C) Network representation of all clusters with size ≥ 5 at the time of maximal cluster size in the trajectory shown in panel B. Each node represents one mRNA molecule and each edge represents one realized intermolecular hybridization, revealing a sparse, branched topology. (D) Cumulative fraction of total mRNA mass contained in clusters up to a given size at steady state. On average, ~60% of total mRNA mass is present in dimers or larger assemblies, ~10% in clusters of size ≥ 60, and ~5% in clusters of size ≥ 190. (E) Steady-state cluster-size distribution. Gray circles show the mean number of clusters of a given size observed per frame at steady state. The red dashed line shows that the mean cluster size can be approximate by a power law with exponent ≈ −2.5, consistent with the Flory-Stockmayer mean-field expectation near gelation. (F) Dependence of the size of the largest cluster (solid line) and the total number of intermolecular interactions (dotted line) on the maximum allowed transcript valency. The sharp increase above a valency threshold of ~6 indicates that a small number of highly multivalent, hub-like transcripts disproportionately stabilize large assemblies by linking smaller subclusters. Vertical dashed line marks the threshold at valency = 6.

These assemblies spanned a broad range of sizes. Most were dimers or small oligomers, but a substantial fraction of transcripts coalesced into much larger clusters, with the largest transiently reaching ~900 molecules (Fig. 2B-C). Because longer transcripts were preferentially incorporated, a larger fraction of total RNA mass (~60%) than molecules (~40%) participated in clusters at steady state. On average, roughly 10% of the mRNA mass was part of clusters containing 60 or more molecules, and 5% was part of clusters of ~190 or more (Fig. 2D). Strikingly, clusters of many different sizes coexisted with no preferred size (Fig. 2B-C). The mean cluster-size distribution was well approximated by a power law with an exponent of –2.5 (Fig. 2E), the value expected from Flory-Stockmayer theory for a system near the critical gelation threshold (39). The native transcriptome therefore appears poised near the boundary between a dispersed state and macroscopic network formation.

Despite their size, these assemblies were highly dynamic. Individual duplexes formed and dissolved continuously, and the size of the largest cluster fluctuated rapidly over time (Fig. 2B). Large clusters (size ≳ 700 molecules) typically lost most of their constituents within tens of milliseconds after reaching a local maximum size (Fig. S4). Thus, even in this RNA-only regime, self-association does not produce a static precipitated state. Instead, it yields a rapidly remodeling network of transient intermolecular contacts.

The topology of these assemblies was also informative. Within large clusters, the total number of interactions was comparable to the total number of molecules (Fig. S5). This is characteristic of a branched, tree-like network (Fig. 2C) rather than an extensively crosslinked structure (40). The number of simultaneous interactions per mRNA followed an approximately exponential distribution, with most molecules engaged in only one or two contacts at a time, whereas a small number behaved as highly connected nodes.

This branched topology suggested that large assemblies are effectively collections of small clusters held together by a few highly multivalent hubs (Fig. 2C, Fig. S5). To test this hypothesis, we selectively removed molecules with high valency (valency > 6) from the largest cluster (Fig. 4F). These high-valency hubs accounted for ~4% of all contacts (~100 total interactions). Yet removing them fragmented the main cluster from ~900 molecules into smaller assemblies, the largest of which contained only ~270 molecules. These hubs are typically long transcripts consistent with our observation that longer mRNAs expose more and longer accessible sites (Fig. 1D, E). For example, *gltB*, one of the longest *E. coli* transcripts (4,461 nucleotides), engaged in up to nine simultaneous interactions. Together, these simulations show that RNA chemistry alone can place the native transcriptome near a gelation transition, producing extensive yet highly dynamic self-association organized around long, multivalent mRNAs that act as hubs.

### 3. Purified mRNA aggregates in vitro in agreement with model predictions

If the clustering predicted *in silico* (Fig. 2) reflects an intrinsic property of the mRNA pool, it should become evident when normal cellular constraints are removed. We therefore examined purified *E. coli* mRNA *in vitro*, in the absence of ribosomes, RNA-binding proteins, and other active cellular factors. Purified *E. coli* mRNA readily aggregated *in vitro* (Fig. S6) in the presence of polyethylene glycol (PEG), which mimics intracellular crowding (Fig. 3A). Robust aggregation was already evident at 5 ng/μL RNA, corresponding to ~2% of the estimated intracellular mRNA concentration. At 10% PEG, nearly all RNA partitioned into the aggregated fraction (Fig. S6).

**Figure 3.**
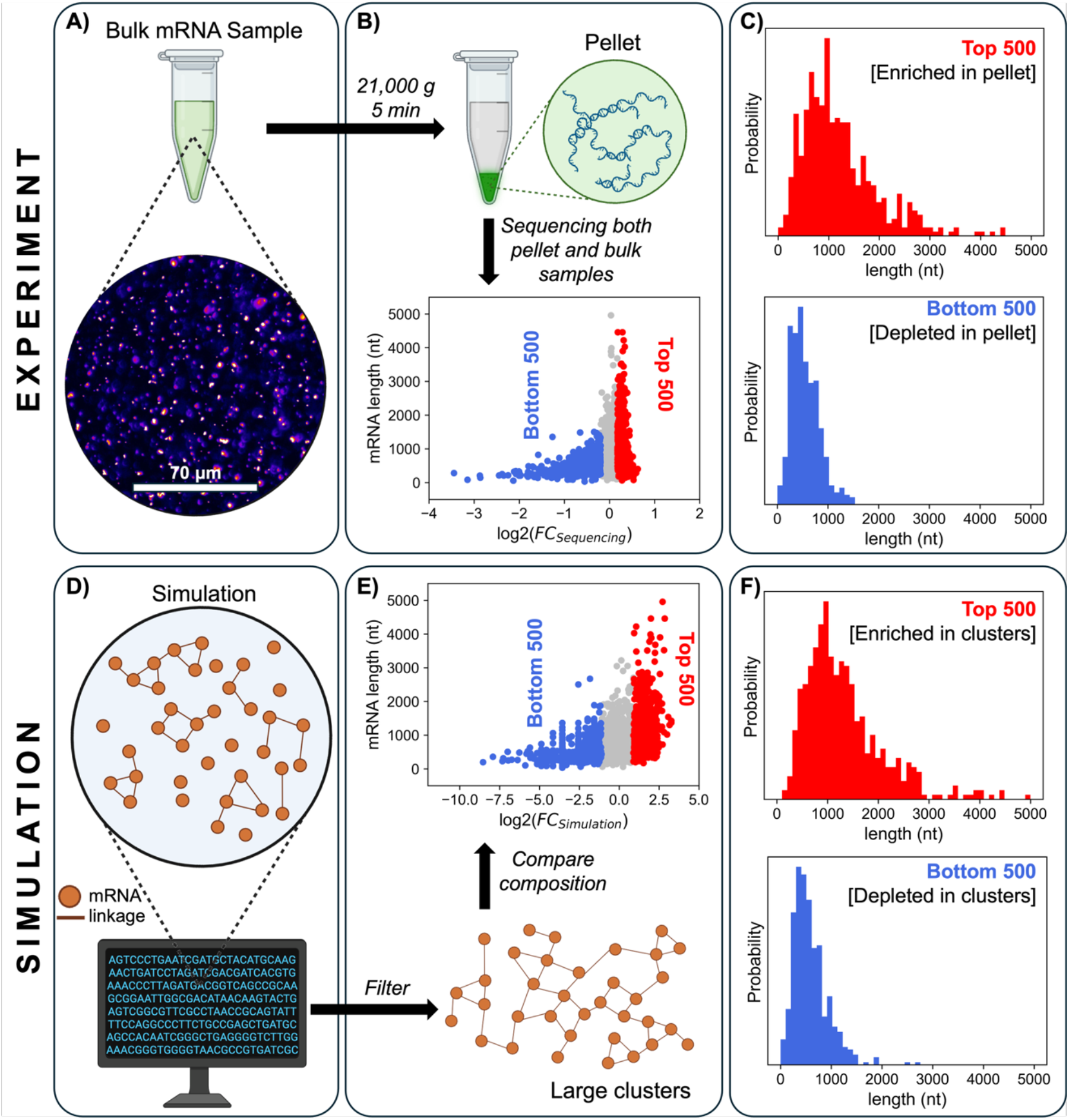
Purified *E. coli* mRNA aggregates *in vitro* with composition matching simulation predictions. (A) Purified *E. coli* mRNA readily aggregates *in vitro*. Representative fluorescence micrograph of mRNA stained with SYBR Gold (5 ng/μL RNA, 10% PEG, 1 M NaCl). (B) Log_2_ fold change (FC) for each transcript, defined as its abundance in the aggregated pellet relative to the input, plotted as a function of transcript length. Positive FC indicates enrichment in the aggregated fraction and negative FC indicates depletion. The top 500 enriched and bottom 500 depleted transcripts are highlighted. (C) Length distributions of the top and bottom 500 transcripts ranked by experimental FC, showing preferential enrichment of long transcripts in the aggregated fraction. (D) Representative snapshot from the simulation illustrating formation of large mRNA clusters in the RNA-only model. (E) Simulation-derived log_2_ FC for each transcript, defined as its relative abundance in the largest cluster compared with its abundance in the full transcriptome, plotted as a function of transcript length. The top and bottom 500 transcripts by FC are highlighted. (F) Length distributions of the top and bottom 500 transcripts ranked by simulation FC, showing the same preferential enrichment of long transcripts in clusters as observed experimentally.

To test whether this behavior required RNA-RNA interactions rather than crowding alone, we examined homopolymeric RNAs under comparable conditions (Fig. S7). PolyU and polyC did not form detectable aggregates, whereas polyA and polyG readily aggregated under the same conditions, in line with previous reports that these homopolymers can self-associate (41, 42). Mixtures of complementary homopolymers also formed aggregates (Fig. S7). These experiments indicate that molecular crowding alone is insufficient to drive aggregation and that attractive inter-molecular base-pairing and/or stacking interactions are required.

These aggregates (Fig. 3A) were sufficiently large to be isolated by differential centrifugation, allowing direct examination of their composition (Fig. 3B, Methods G). We sequenced both the aggregated (pellet) fraction and the total input RNA, and computed a fold change (FC) for each transcript (Fig. 3B). In parallel, we scored each transcript in our simulations by its enrichment in the largest cluster relative to its abundance in the full transcriptome (see Supp. Spreadsheet 4).

The experimental data closely mirrored the simulations (Fig. 3D, E). In both cases, long transcripts were preferentially enriched in the aggregated fraction, while shorter transcripts were depleted. The 500 most enriched transcripts had a median length of ~1,100 nt in both settings, whereas the 500 most depleted had a median length of ~500 nt (Fig. 3C-F). Transcript length alone predicted partitioning with high accuracy (area under the receiver operating characteristic curve (AUC) for simulation = 0.84; for experiment = 0.82; Fig. S8), indicating that the same length-dependent principles that organize clustering *in silico* also shape aggregation *in vitro*. This agreement between experiments and simulations was also evident at the level of individual mRNA species (Fig. S9). Among the 500 most enriched transcripts, 186 were shared between the *in vitro* and *in silico* datasets, far exceeding chance (*p*-value = 1.7 × 10^−8^), while 209 were shared among the two depleted sets (*p*-value = 2.1 × 10^−16^) (Fig. S9).

Together, these results show that our RNA-only model captures an intrinsic physicochemical property of the *E. coli* transcriptome. When cellular constraints are removed, native mRNAs aggregate (Fig. 3A) in a sequence- (Fig. S9) and length-dependent manner (Fig. 3C) that closely matches simulation predictions (Fig. 3F). This agreement suggests that self-association is a latent property of the native mRNA pool.

### 4. The native transcriptome resists aggregation relative to randomized alternatives

Is the self-association we observe simply an unavoidable consequence of any large RNA pool, or have native mRNA sequences evolved to limit it? To test this, we compared the native *E. coli* transcriptome to two classes of randomized controls (Fig. 4A). In the first, codons were shuffled synonymously, so that each transcript encoded the same protein but with a different nucleotide sequence. In the second, dinucleotide frequencies were preserved while all higher-order sequence information was randomized. Dinucleotide shuffling is a particularly stringent control because it conserves nearest-neighbor stacking energies (20), the primary determinant of RNA base-pairing stability. For each class, we generated three independent alternative transcriptomes, maintaining the native transcript lengths and copy numbers, and subjected them to the same simulation pipeline.

**Figure 4.**
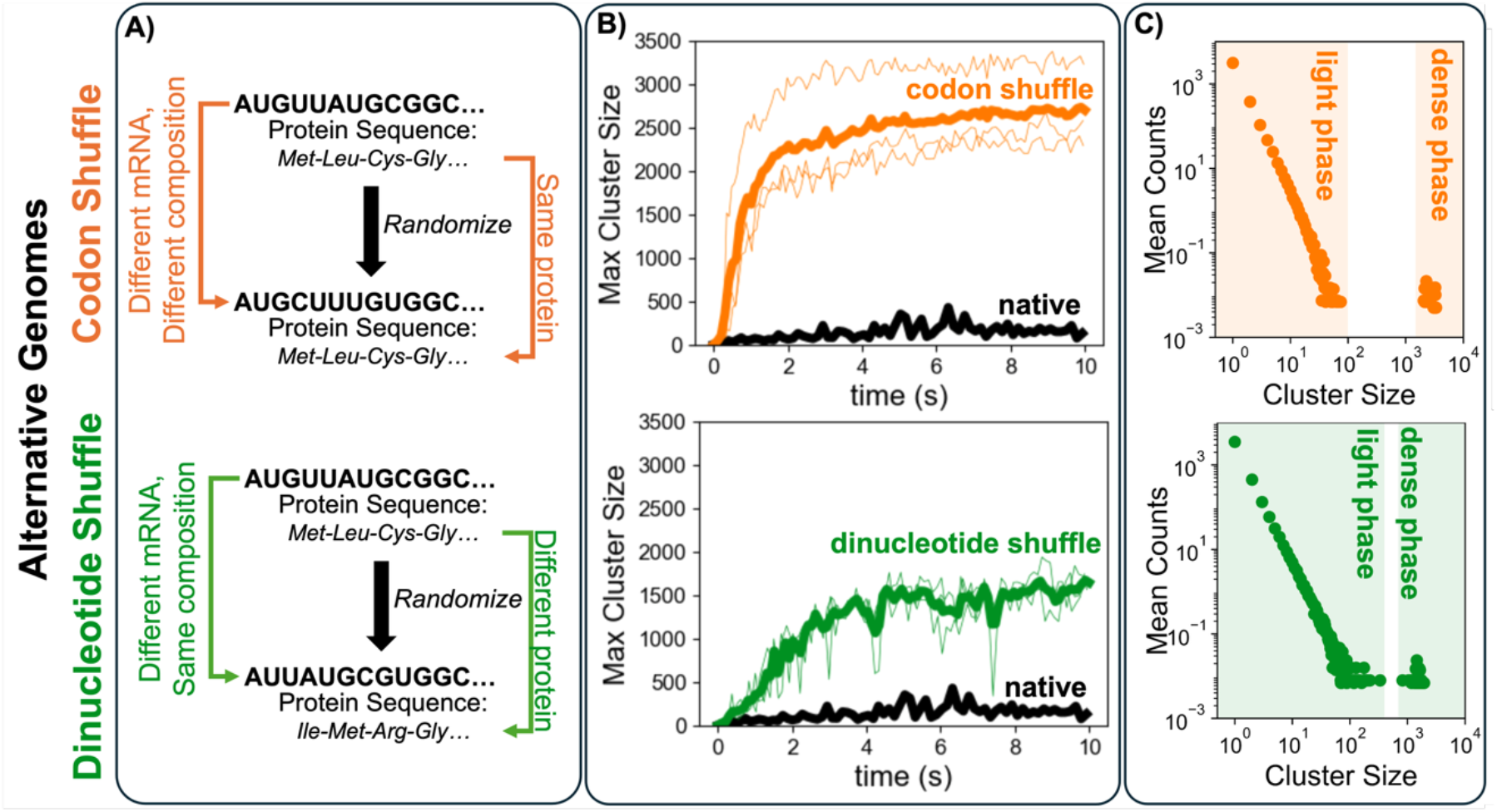
The native *E. coli* transcriptome resists aggregation relative to randomized controls. **(A)** Two classes of randomized control transcriptomes were generated: codon-shuffled (synonymous codons permuted, preserving encoded proteins), and dinucleotide-shuffled (dinucleotide frequencies were preserved while higher-order sequence information was randomized). Three independent transcriptomes were generated for each control class while maintaining native transcript lengths and abundances. **(B)** Size of the largest cluster over time for native (black), codon-shuffled (orange) and dinucleotide-shuffled (green) transcriptomes. Thin lines, individual replicates (N = 3); thick lines, mean. Randomized transcriptomes form markedly larger assemblies than the native transcriptome. **(C)** Steady-state cluster-size distributions for the randomized transcriptomes. The native transcriptome follows a continuous power-law distribution (see Fig. 2E), whereas both randomized controls exhibit bimodal distributions with many small clusters and a distinct population of very large assemblies, consistent with the emergence of a percolated, gel-like network. These large connected assemblies account for ~40% of total mRNA mass in codon-shuffled transcriptomes and ~30% in dinucleotide-shuffled transcriptomes.

All randomized transcriptomes were markedly more prone to aggregation than the native transcriptome (Fig. 4B). The largest clusters in codon-shuffled and dinucleotide-shuffled simulations contained as many as ~3,300 and ~1,500 molecules, respectively, compared to ~900 in the native transcriptome. The native and randomized transcriptomes differed not only in the extent of clustering but in the regime of self-association. In the native transcriptome, there is no preferential cluster size; the size of clusters is continually distributed following a power-law as expected near a gelation transition (Fig. 2E). In contrast, both codon-shuffled and dinucleotide-shuffled controls, by contrast, produced a bimodal cluster-size distribution: many small clusters coexisted with a second population of very large assemblies, with relatively few intermediate-sized clusters (Fig. 4C). This marks a clear shift away from the near-critical regime of the native transcriptome and is consistent with the system crossing a gelation threshold. In this state, large interconnected assemblies emerge and capture a substantial fraction of total mRNA pool (~40% of total mRNA mass in codon-shuffled and ~30% in dinucleotide-shuffled transcriptomes), despite identical transcript abundances across all simulations.

These results demonstrate that the native *E. coli* transcriptome, while intrinsically prone to aggregation, is far less so than matched randomized alternatives. Because the codon-shuffled controls preserve protein output and the dinucleotide-shuffled controls preserve local base-stacking composition, this difference cannot be explained simply by coding constraints, transcript length, abundance, or bulk nucleotide composition. Instead, it suggests that natural selection has shaped mRNA sequences to limit promiscuous intermolecular pairing and avoid the extensive clustering seen in randomized transcriptomes.

### 5. Native mRNA sequences bear signatures of selection against promiscuous intermolecular interactions

Having established that the native *E. coli* transcriptome is markedly less prone to self-association than matched randomized alternatives (Fig. 4), we next asked which sequence-level properties underlie this difference. We focused on the top 100 most abundant mRNAs in *E. coli*. Together, these mRNAs account for ~3,600 molecules at steady state and participate in ~1 million potential interactions in our simulations (see Supp. Spreadsheet 3). For each transcript, we generated 100 matched variants using both synonymous codon shuffling and dinucleotide-preserving shuffling. We then compared native and randomized sequences using three complementary metrics: intramolecular folding stability, accessible single-stranded sequence, and the strength of the most favorable intermolecular interaction that the transcript can form with other mRNAs in the transcriptome (Fig. 5, Methods J; Supplementary Note 4; Fig. S10).

**Figure 5.**
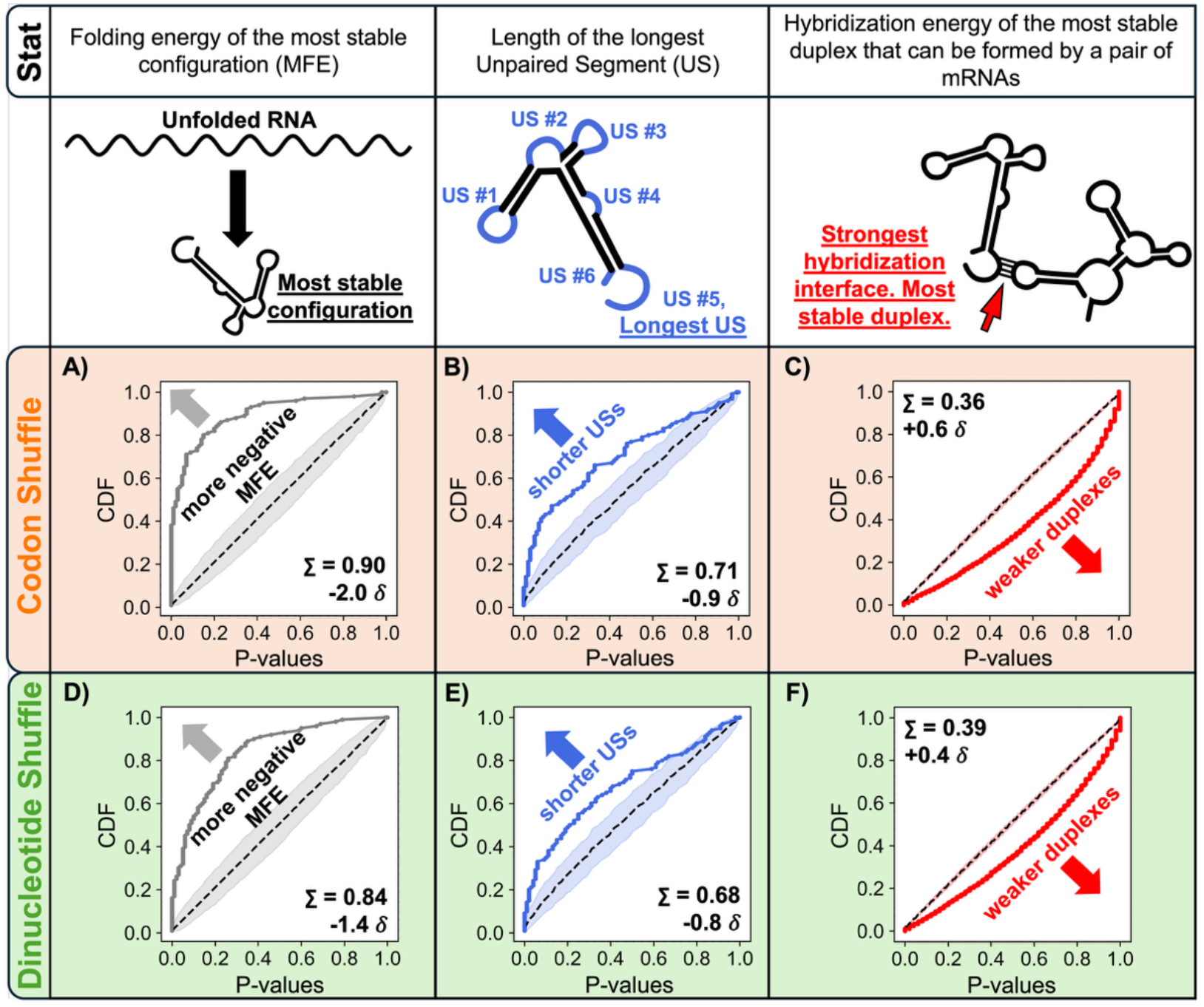
Native *E. coli* mRNAs fold more stably, expose shorter unpaired segments, and form weaker intermolecular duplexes than randomized controls. The top row defines the three statistics analyzed: minimum free energy (MFE) of the most stable intramolecular fold, length of the longest unpaired segment (US), and hybridization energy of the most stable intermolecular duplex predicted for a pair of mRNAs. (A-C) Comparison with codon-shuffled sequences. (D-F) Comparison with dinucleotide-shuffled sequences. Each panel shows the cumulative distribution of one-tailed P-values for the indicated metric, where each P-value represents the fraction of shuffled variants with a value at least as low as the corresponding native sequence (see Supplementary Note 4 and Fig. S10). Dashed line indicates the null expectation (Σ ≈ 0.5), and shaded region shows the 95% confidence interval. Σ quantifies the area under the CDF and ranges from 0 to 1. For folding energy (A, D), Σ > 0.5 indicates that native sequences consistently fold more stably than shuffled controls; for USs (B, E), Σ > 0.5 indicates that the longest unpaired segment in native sequences is consistently shorter than shuffled controls; for intermolecular duplex strength (C, F), Σ below 0.5 indicates that native sequences form weaker duplexes. δ expresses the magnitude of the difference in standard deviations of a matched normal distribution. See Supplementary Note 4 for full statistical framework.

To compare these metrics between native and shuffled sequences, we used a non-parametric statistical framework (Supplementary Note 4, Fig. S10). For each gene, and each intramolecular metric, we computed the one-tailed probability that a shuffled variant would yield a value at least as low as the native sequence (e.g., more negative folding free energy). For the intermolecular metric, the same analysis was performed across transcript pairs. These one-tailed P-values were then pooled and summarized by the area under their cumulative distribution (∑). Under the null expectation of no systematic difference between native and shuffled sequences, P-values are uniformly distributed and ∑ ≈ 0.5. Systematic deviations from 0.5 therefore indicate consistent differences between native and randomized sequences, with ∑ close to 1.0 reporting on the tendency of P-values to be small, and ∑ close to 0.0 reporting on P-values being systematically large. To aid interpretation, we also report an equivalent effect size,Σ, defined as the shift in standard deviation units of a matched normal distribution that reproduces the observed Σ (Supplementary Note 4, Fig. S10).

Native mRNAs consistently achieved more stable intramolecular folding energies than their randomized counterparts (Fig. 5A, D, Fig. 6A, Supplementary Note 5). More stable folding would sequester a greater fraction of the sequence in intramolecular structure, thereby reducing the number of unpaired bases available for promiscuous intermolecular contacts. To assess whether this effect could arise by chance and to determine whether it was sufficiently large to propagate to transcriptome scale, we performed a bootstrap analysis (Fig. 6, Methods J). We assembled 10,000 random sets, each containing one shuffled variant per gene from the top 100 most abundant mRNAs and compared their mean folding energy to that of the native set. None of the 10,000 codon-shuffled or dinucleotide-shuffled transcriptome sets matched the native mean folding energy (empirical P-value < 10^−4^, Fig. 6B).

**Figure 6.**
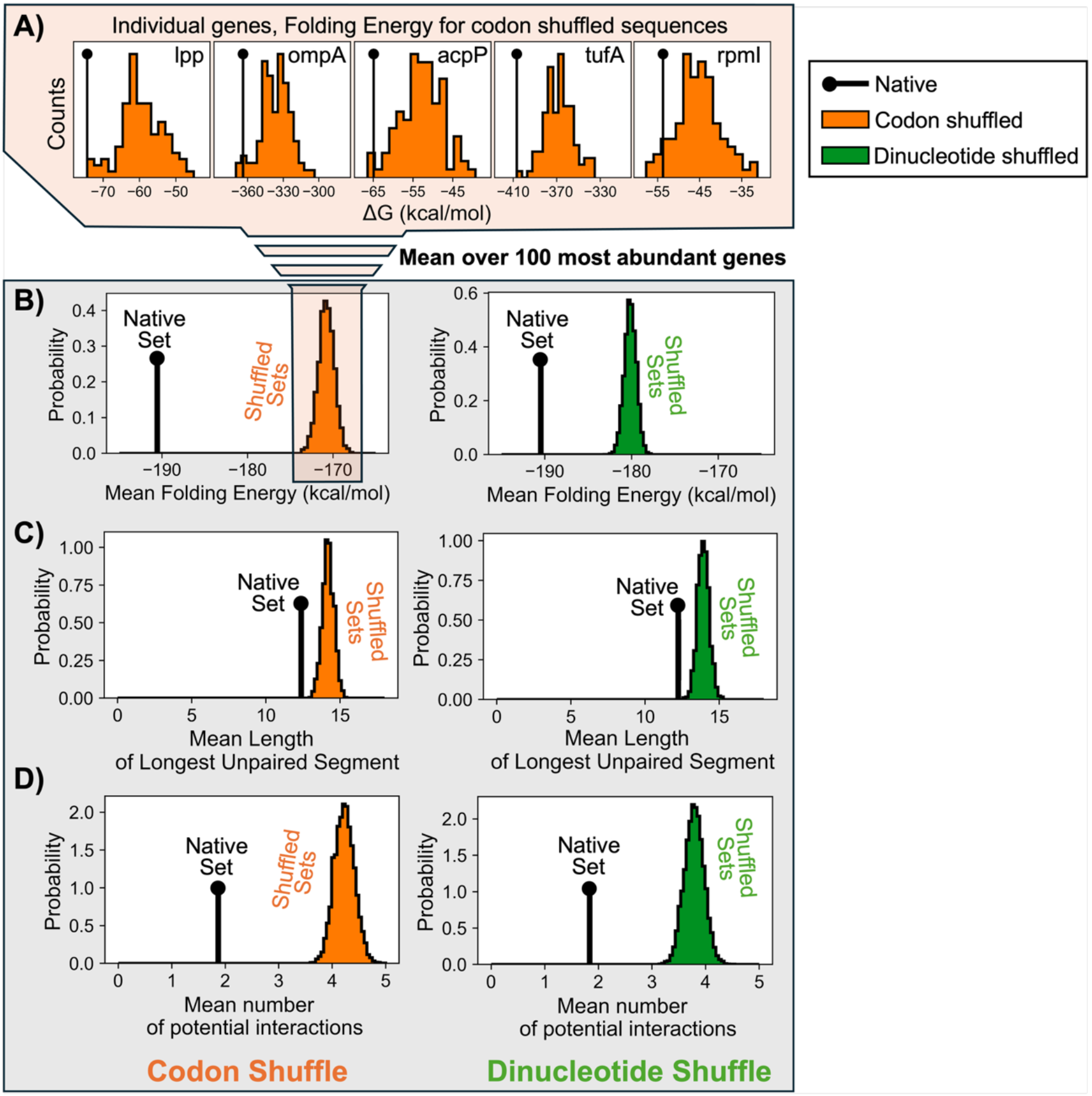
Native *E. coli* mRNAs bear signatures of selection against promiscuous intermolecular pairing. (A) Representative examples from the five most abundant *E. coli* transcripts comparing native sequences (black) with codon-shuffled counterparts (orange) show that native mRNAs generally achieve more stable intramolecular folding energies. (B) Mean intramolecular folding free energy of the native set of the top 100 most abundant transcripts compared with bootstrap distributions generated from 10,000 matched randomized sets assembled from codon-shuffled variants (left) or dinucleotide-preserving shuffled variants (right, in green). Native sequences are more stably folded than all randomized sets. (C) Mean length of the longest unpaired single-stranded segment in the native set compared with the corresponding bootstrap distributions from codon-shuffled and dinucleotide-shuffled controls. Native sequences expose shorter accessible regions than randomized controls. (D) Mean number of strong candidate intermolecular interaction partners per transcript in the native set compared with the corresponding randomized controls. Strong candidate interactions were defined using the most favorable duplex for each transcript pair and a threshold of ΔG ≤ −7 kcal/mol. Native sequences harbor fewer strong intermolecular contacts than randomized controls.

Native sequences also exposed shorter single-stranded regions upon folding. In particular, the longest accessible single-stranded segment in each transcript was shorter in native sequences than in the shuffled controls (Fig. 5B, E). Again, none of the 10,000 random sets matched the native transcriptome in this metric (Fig. 6C), regardless of the randomization scheme. These trends were observed broadly across the abundant mRNA set rather than being driven by a few outliers. Thus, native coding sequences are biased toward structures that both sequester more RNA internally and leave less exposed sequence available for promiscuous pairing.

We next asked whether these structural differences translated into weaker promiscuous intermolecular pairing. For each mRNA pair, we determined the hybridization energy for the most favorable duplex and compared native sequences with their matched randomized counterparts (Fig. 5C, F). Native sequences consistently formed weaker intermolecular interactions than either codon-shuffled or dinucleotide-shuffled controls (Fig. 6D). Randomized sets also harbored disproportionately more strong candidate interactions (*ΔG*_*min*_ < −7 kcal/mol) than the native set (Fig. 6D), providing a direct explanation for the increased aggregation observed in the transcriptome-wide simulations (Fig. 4). Taken together, these results demonstrate that the reduced self-association of the native transcriptome is not a chance outcome of coding constraints or nucleotide composition.

To test whether these sequence signatures extend beyond bacteria, we performed the same analysis on a set of 10 highly abundant human mRNAs (Methods J). Consistent with the trends observed in *E. coli*, native human sequences tended to fold more stably and expose shorter unpaired regions than codon-shuffled controls (Fig. 7). These results suggest that selection against promiscuous intermolecular pairing extends beyond bacterial transcriptomes.

**Figure 7.**
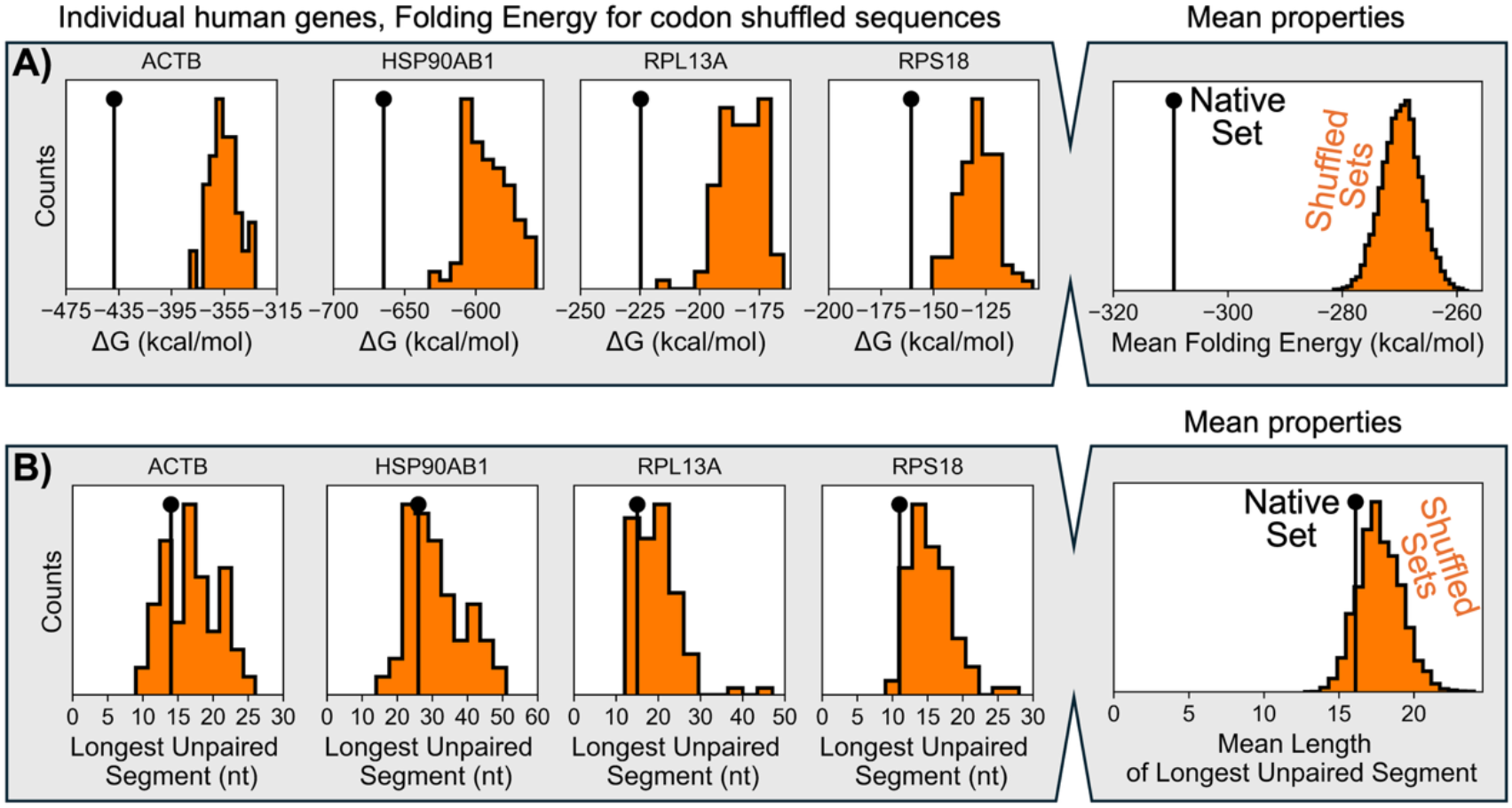
Human mRNAs show similar signatures of selection against promiscuous intermolecular pairing. Representative examples from four abundant human transcripts comparing native sequences (black) with codon-shuffled counterparts (orange). (A) Native mRNAs achieve more stable intramolecular folding energies, and (B) expose shorter longest accessible single-stranded segments than shuffled controls. Right panels show the mean properties for a set of 10 abundant mRNAs compared to the bootstrap distributions generated from 10,000 matched randomized sets.

These analyses reveal pervasive signatures of selection against promiscuous intermolecular RNA pairing in native transcriptomes. Native mRNAs are biased toward stronger intramolecular folding, shorter accessible single-stranded regions, and weaker intermolecular complementarity than matched randomized controls. Together, these features argue that transcriptome solubility has been a direct target of natural selection, encoded in mRNA sequence alongside protein-coding information.

## Discussion

Our findings identify transcriptome self-association as an intrinsic consequence of RNA chemistry. When considered as a population rather than as isolated molecules, mRNAs are not simply carriers of coding information. Their base-pairing chemistry, combined at transcriptome-scale concentrations, creates a latent capacity for extensive intermolecular association and higher-order assembly. In an RNA-only setting, transcriptome-scale simulations predict widespread, dynamic clustering (Fig. 2), and purified mRNA recapitulates this behavior *in vitro* (Fig. 3). The observation that native sequences are depleted of features that promote promiscuous intermolecular pairing (Fig. 5-7) indicates that this pressure is not merely physically plausible, but has shaped genome composition.

This framework helps reconcile two otherwise conflicting observations. RNA aggregates readily *in vitro*, yet widespread mRNA aggregation is not observed in unperturbed cells. Our results suggest that this discrepancy reflects not the absence of an underlying physical drive but its active suppression. Translation and ribosome occupancy are likely central to this buffering, masking mRNA regions that would otherwise remain available for intermolecular contact. RNA-binding proteins, helicases, and turnover may further suppress or resolve promiscuous RNA interactions. The RNA-only baseline defined here therefore provides a quantitative reference for assessing how much buffering is required to maintain mRNAs dispersed *in vivo*.

An important implication is that self-association is a collective property of the transcriptome, not a feature of individual molecules. The folding propensity, accessible sequence, and intermolecular complementarity of any single mRNA may appear modest in isolation. But these features compound across thousands of coexisting species in a confined cellular volume, and together they determine the physical state of the mRNA pool as a whole. Consistent with this view, the native transcriptome occupies a near-critical regime whose cluster-size distribution is close to the Flory-Stockmayer mean-field expectation for gelation, whereas randomized transcriptomes cross this threshold, producing giant connected assemblies.

These results also highlight the evolutionary significance of RNA structure in coding sequences. Earlier sequence-shuffling studies suggested that structural features of native coding sequences may differ from randomized controls, although the interpretation of those differences remained unsettled (43–47). mRNA sequences in bacteria and archaea have evolved to avoid stochastic interactions with abundant ncRNAs such as rRNAs and tRNAs (48). Our results extend that logic to the coding transcriptome itself. Because mRNAs are long, partially unstructured, and collectively present at high concentration, the combinatorial space for promiscuous contacts is far larger than for short, stably folded noncoding RNAs, making this a quantitatively distinct and potentially more demanding constraint on sequence evolution.

Although the quantitative framework developed here was built around *E. coli*, the underlying logic should not be unique to bacteria. The ingredients that generate accidental RNA-RNA complementarity: a limited alphabet, high molecular abundance, finite volume, and incomplete intramolecular sequestration, are general features of all cells. The similar signatures we observe in abundant human mRNAs (Fig. 7) support this view. This constraint may in fact be more acute in eukaryotes, where pre-mRNAs must fold and undergo processing in the nucleus before engaging the translation machinery, leaving nascent RNA vulnerable to promiscuous intermolecular pairing.

These evolutionary signatures also bear on codon choice. Synonymous codons are not freely interchangeable if alternative coding sequences alter structural features that affect folding stability and unintended intermolecular pairing. This constraint operates on the same sequences that must also satisfy protein-coding requirements, translational kinetics, and regulatory signals. Transcriptome solubility therefore represents an additional layer of constraint on coding-sequence design, operating alongside, and at times potentially in tension with translational optimization. Consistent with this, recent work has shown that mRNA secondary structure can influence protein output through changes in functional half-life, independent of codon optimality (49).

Our model is intentionally reductionist. It omits ribosomes, RNA-binding proteins, helicases, and active turnover, and relies on thermodynamic parameters derived largely from *in vitro* measurements on simpler systems. RNA behavior in cells may differ from dilute-solution expectations. Recent work suggests that the intracellular environment may modestly weaken base-pairing (50), and large-scale *in vivo* probing has shown that mRNAs are often less structured in cells than *in vitro* (51, 52). At the same time, molecular crowding in cells could favor repeated collisions and stabilize RNA hybridization (53, 54). Despite these simplifications, the agreement between our simulations and purified-mRNA experiments (Fig. 3, Fig. S9) indicates that the model captures a real and consequential property of the mRNA pool. The remaining gap between the self-association predicted by our RNA-only baseline and the relative dispersal observed in living cells points either to strong buffering mechanisms or to aspects of intracellular RNA thermodynamics that remain poorly understood (55).

More broadly, our results suggest that cells must continually solve a transcriptome-solubility problem. This framework yields testable predictions: self-association pressure should rise when the free mRNA pool increases, such as during polysome disassembly, or when conditions favor stabilization of non-productive duplexes as may occur during cold stress. The dramatic upregulation of cold shock proteins like CspA in *E. coli*, specialized RNA chaperones that resolve inappropriate duplexes (56), is consistent with this view. From this perspective, many aspects of RNA metabolism such as co-transcriptional folding (57), surveillance by RBPs (30), and rapid mRNA turnover (58, 59), can be viewed, at least in part, as solutions to the problem of keeping transcriptomes soluble. The same multivalent interactions that become deleterious when unchecked may also contribute to functional RNA-rich condensates, where long RNAs are often preferentially enriched (9). By defining an RNA-only reference state and showing that native sequences are biased away from extensive self-association (Fig. 4-7), this work provides a framework for investigating how cells manage their RNA pools at transcriptome scale.

## Materials and Methods

### A. Sequencing data and RNA abundance for simulation

To study the *E. coli* transcriptome, we used the K-12 MG1655 genome (assembly ASM584v2) obtained from RefSeq (60). RNA sequences were taken from the recently published compendium from Tjaden (34) (available in the Harvard Dataverse at https://doi.org/10.7910/DVN/QBMC9D, Assemblies Table 2) and ncRNA, rRNA and tRNA were filtered out. For each mRNA, a median transcript abundance was computed and stored in an array *A*.

Assuming 7,800 mRNA molecules in a single *E. coli* cell grown in LB (32) we converted the copy number for the *i*-th mRNA based on its transcript abundance as:

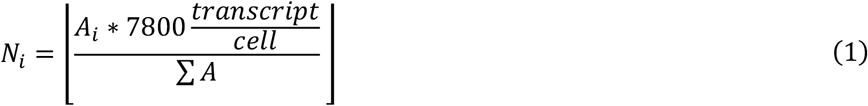

This yielded a total of 7,569 RNA molecules after rounding down.

### B. RNA folding *in silico*

To determine accessible regions along the transcripts that are potentially available for base-pairing, the RNA sequences were folded *in silico* using the ViennaRNA 2.6.4 Python library on the Whitehead Institute BaRC Fry cluster. Instead of relying on the minimum free energy (MFE) structure, we opted for a more conservative approach previously established for the prediction of antisense oligonucleotides binding sites (61), where every nucleotide *i* over one mRNA of length *L* is considered as “weakly paired” when:

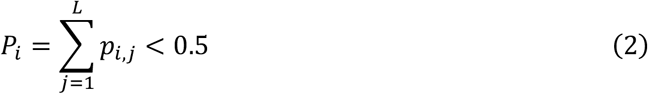

where *p*_*i,j*_ is the base-pair probability for bases *i* and *j*. When compared to directly using the centroid structure of the Boltzmann ensemble (62), our approach is more stringent, although as a downside it does not necessarily return a single physically meaningful fold. Because *in silico* predictions tend to overestimate structure (51, 52, 63), this approach likely underrepresents the true number of accessible regions and serves as a conservative estimate of interaction potential (see Supplementary Note 6 and Fig. S11 for evaluation of ViennaRNA performance).

### C. Prediction of interactions among *E. coli* mRNAs

To evaluate potential interactions among mRNAs, we first retrieved single stranded regions along the transcripts. To do so, for each mRNA, we extracted stretches having at least 7 adjacent “weakly paired” nucleotides (nts). The 7-nt threshold was chosen to avoid introducing an excessively large number of extremely short stretches that would have made computations unfeasible. Importantly, in this work we disregarded the role of RNA strand displacement to be more conservative (24, 27), and discarded stretches that would have been 7 or more nt-long if they were not interrupted by just one or two “strongly paired” nucleotides.

For every pair of transcripts, we computed the MFE for hybridization between any of their annotated unstructured segments using ViennaRNA *duplexfold* function. A triangular (7569 x 7569) interaction matrix was filled at any *i,j* entry by storing the most stable hybridization energy among all possible hybridizing interfaces for any two transcripts *i* ad *j* (*ΔG*_*min*(*i,i*)_). For each entry of the matrix, a corresponding equilibrium constant was calculated:

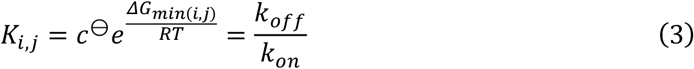

with T being the absolute temperature (310.15 K), R the gas constant (1.987 × 10^−3^ kcal/mol/K) and c_⊖_ the standard reference concentration (1 mol/L).

### D. Kinetic stochastic simulations

To evaluate the RNA interaction potential inside an *E. coli* cell, we set up a custom kinetic stochastic simulation. The code was written in Python and accelerated using just-in-time (JIT) compilation provided by the Numba library (64), and executed on the Whitehead Institute BaRC Fry cluster.

The simulation strategy was based on the Gillespie algorithm for the simulation of networks of chemical reactions (38). Briefly, at each time point *t* of the simulation, the state of the system is defined by the comprehensive list of all RNA molecules and their state (paired/unpaired). From this list, we compute “on the fly” all possible reactions *i ∈ χ* that can occur in the current state, and their respective rates *r*_*i*_. To evolve the system, we (i) sample the waiting time *τ* for the next reaction from the exponential distribution with mean equal to 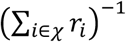 to update *t → t + τ*, and (ii) randomly select a reaction by sampling with probability proportional to 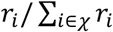. The two steps are repeated until the simulation reaches the chosen time limit, and the state of the system is exported at fixed intervals for further analysis (see Supplementary Note 7 for additional details).

### E. Chemicals

The following reagents were used in this work: polyA (Sigma Aldrich, P9403), polyU (Sigma Aldrich, P9528), polyC (Sigma Aldrich, P9403), polyG (Sigma Aldrich, P4404), 5M NaCl solution (Invitrogen, AM9760G), Polyethylene Glycol 8K (Sigma Aldrich, P5413-1KG), SYBR Gold (Thermo Scientific, S11494), Phenol:Chloroform: Isoamyl Alcohol, 25:24:1, Saturated with 10mM Tris, pH 8.0, 1mM EDTA (Sigma Aldrich, P3803-100ML), Lysozyme (Thermo Scientific, 89833), Phosphate Buffered Solution, pH 7.4 (PBS; Gibco, 14190250), 1M Tris pH 8.0 (Invitrogen, AM9855G), 0.5M EDTA pH 8.0 (Invitrogen, AM926), 3M Sodium Acetate pH 5.5 (Invitrogen, AM9740), Nuclease-Free Water (Qiagen, 129114), Ethyl Alcohol 200 Proof (Ethanol; Pharmco, 111000200), Isopropanol, 99.5% (Thermo Scientific Chemicals, AC383920025), TURBO DNase™ (2 U/μL) (Thermo Fisher Scientific, AM2238), AMPure XP Beads for DNA Cleanup (Beckman, A63881) and MICROBExpress™ Bacterial mRNA Enrichment Kit (Invitrogen, AM1905).

### F. *E. coli* RNA extraction and mRNA enrichment

5 milliliters of LB was inoculated with *E. coli* K-12 MG1655, GLB 116, and grown overnight at 37 °C. The following day, the overnight culture was backdiluted 1:100 in LB and resuspended. This back-diluted culture was then diluted to a final concentration of 1:10,000 in 15 mL of LB in a 125 mL Erlenmeyer flask (Cole-Parmer Essentials Boro 3.3 34502-58). This flask was placed in a 37°C shaker room and allowed to grow in exponential phase for 10 doublings before harvest, approximately 4 hours and 10 minutes. The OD600 was then measured at 0.244 using the Ultrospec 2100 pro (Amersham Biosciences). The flask was decanted into a 50mL falcon tube (Falcon, 352070) and spun down at 4°C and 4000 g for 10 minutes. The LB was removed, and the pellet was resuspended in 200 uL of PBS (Gibco, 14190250) and placed in a −80 °C until use.

The RNA extraction method was adapted from Gill *et al* (65). After spinning down the thawed pellet, the supernatant was removed and the pellet was resuspended in 10 mg/mL Lysozyme prepared in TE Buffer (10mM Tris, pH 8, 1mM EDTA) and incubated at room temperature for 5 minutes. The total RNA was extracted by phenol:chloroform, precipitated and washed with isopropanol and ethanol. After extraction, the total RNA was further purified using TURBO DNase™ (2 U/μL) and cleaned using AMPureXP Beads at 2.0x concentration of initial reaction volume.

The MICROBExpress™ Bacterial mRNA Enrichment Kit was used with minimal modifications: all precipitations were performed overnight at −80°C and spun down at 16,000g for 20 minutes at 4°C, the second wash of the OligoMagBeads was collected separately from the first elution and the resulting mRNA was resuspended in Nuclease-Free Water.

### G. Aggregation of *E. coli* mRNA *in vitro*

To evaluate whether the mRNA extracted from *E. coli* was poised to form aggregates, we prepared samples for microscopy observation by mixing variable amounts of mRNA, PEG 8k, NaCl and SYBR Gold in Nuclease-Free Water. The samples prepared this way were briefly annealed by heating them up to 60°C and cooling them down to room temperature to reduce kinetically stuck configurations. Observations of the aggregates were performed on an Invitrogen Evos 7000 inverted microscope.

The samples prepared for sequencing were obtained by mixing 25 ng/μL of mRNA, 6.5% PEG 8k, 1 M NaCl, 1x SYBR Gold in a 500 uL Eppendorf tube. The aggregates were pelleted by centrifugation at 21,130g for 5 minutes (Eppendorf Centrifuge 5424). The supernatant was removed and the pellet diluted in Nuclease-Free Water before being annealed to solubilize the aggregates. The concentration of the resuspended pellet was measured using a Nanodrop 8000 (Thermo Scientific), which led to an estimate of ~40% of the total mRNA mass being in the aggregated fraction.

### H. Library preparation and sequencing

Directional RNA-seq libraries were constructed by Novogene using random hexamer priming for first-strand cDNA synthesis, followed by second-strand synthesis incorporating dUTP to preserve strand specificity. Libraries were end-repaired, A-tailed, adapter-ligated, size-selected, USER-treated, PCR-amplified, and purified according to standard Illumina protocols. Library quality and concentration were assessed by Qubit fluorometry, real-time PCR, and Bioanalyzer analysis.

RNA from each of the two samples (input and aggregated fraction) was used to generate three independent library-preparation technical replicates for RNA-seq (six libraries total). These were sequenced on an Illumina NovaSeq X Plus platform, generating paired-end reads with a target output of 3.0 G per library. Raw reads were subjected to quality filtering by the sequencing provider to remove adapter contamination, reads containing >10% undetermined bases, and reads with >50% low-quality bases (Q ≤ 5). The resulting high-quality reads were used for all downstream analyses.

### I. Sequencing data analysis

An independent quality-control step was performed to ensure consistent preprocessing across samples. Paired-end RNA-seq reads (2 × 150 bp) were quality-controlled and filtered using *fastp* (66) (v0.23.4), resulting in minimal read loss (>99% reads retained) and negligible changes in read length.

Filtered reads were aligned to the *E. coli* K-12 MG1655 reference genome (assembly ASM584v2) using *Bowtie2* (67) (v2.4.2), and alignments were sorted and indexed with *samtools* (68) (v1.11). Read quantification was performed using *featureCounts* (69) (v2.0.2) assigning paired-end reads to annotated genomic features based on the provided GTF annotation. Only properly paired reads were counted, and reverse-strand specificity was enforced during counting.

Raw read counts were normalized to counts per million (CPM) by dividing gene-level counts by the total number of assigned reads across all features, including rRNA, and multiplying by 10^6^. CPM values were then averaged across the three technical replicate libraries for each sample. Genes with an average CPM greater than 2 in at least one of the two samples were retained for further analysis. Unless otherwise stated, downstream analyses were restricted to protein-coding genes.

To reduce length-bias in sequencing efficiency, transcript abundance was additionally quantified as transcripts per million (TPM) by first normalizing read counts by gene length and subsequently scaling length-normalized counts to a per-million basis across all protein-coding genes. This correction yielded good agreement between transcripts abundance in our dataset and the reference data used to setup the simulation (Fig. S12).

Finally, to quantitatively compare sequencing and simulation data, we kept only the intersecting elements between the subset of highly expressed genes retained in our sequencing dataset (2611) and the subset of genes part of our simulation (1952), for a total of 1815 common entries.

### J. Statistical analysis of shuffled sequence

To test whether native mRNA sequences are biased against promiscuous intermolecular self-association, we analyzed highly abundant transcripts in *E. coli* and human. For *E. coli*, we selected the top 100 most abundant mRNAs from the reference dataset. For each transcript, we generated 100 synonymous codon-shuffled variants and 100 dinucleotide-preserving shuffled variants. Codon shuffling preserves the encoded protein while altering nucleotide sequence, whereas dinucleotide-preserving shuffling preserves the base-pairing potential while disrupting higher-order sequence organization. For human, we analyzed 10 abundant mRNAs (ACTB, B2M, EEF1A1, FTH1, GAPDH, HSP90AB1, RPL13A, RPLP0, RPS18, TPT1) and generated 100 codon-shuffled variants per gene.

For each native and shuffled sequence, we computed metrics related to aggregation propensity: intramolecular folding free energy, the length of the longest accessible single-stranded segment after folding, and, for the *E. coli* set, intermolecular interaction strength derived from the most favorable duplex formed with other abundant mRNAs. Because human transcripts are substantially longer, folding-related quantities for the human analysis were computed using LinearPartition (70) rather than ViennaRNA.

To compare native sequences with their matched shuffled controls in *E. coli*, we used an empirical non-parametric framework described in detail in Fig. S10 and Supplementary Note 4. To assess whether these sequence-level differences propagate to the transcriptome level, we performed a bootstrap analysis by assembling 10,000 random sets, each containing one shuffled variant per gene, and comparing the mean statistic of each set with that of the native transcript set. This analysis was performed for folding energy and longest accessible segment in both *E. coli* and human, and for intermolecular interaction propensity in *E. coli*, summarized as the number of strong candidate intermolecular partners per transcript using a threshold of ΔG ≤ −7 kcal/mol.

## Acknowledgments

This work is supported by grants from the NIH (R35GM151111), Chan Zuckerberg Initiative (DAF2022-250422), Richard and Susan Smith Family Foundation, and the David and Lucile Packard Foundation.

