## Supplementary for "Maintaining transcriptome solubility constrains mRNA sequence composition"

*Corresponding author.

**This PDF file includes:**

Supplementary Notes

Supporting Information

- Figures S1 to S12
- Tables S1 to S2
- SI References

**Supplementary Notes**

| Table of Content | Page |
| --- | --- |
| 1. Indefinite self-association model | 3 |
| 2. Effect of different *k_on_* values on the simulation | 4 |
| 3. Effect of different energy thresholds on the simulations | 5 |
| 4. Sequence-shuffle: analysis pipeline | 6 |
| 5. Sequence-shuffle: results | 7 |
| 6. Quantifying ViennaRNA accuracy | 8 |
| 7. Simulation scheme: additional details | 9 |

1. Indefinite self-association model

To estimate the interacting fraction in a system composed by self-associating molecules, we have implemented a simple indefinite self-association model (equal *K*) as described by Martin (3).

Briefly, for a system of self-associating RNA molecules, it is possible to write a set of successive equilibria and associated equilibrium constants:

$\left[ RNA \right]+\left[ RNA \right]\rightleftharpoons{[RNA]}_{dimer}\Longrightarrow K_{dimer}=\frac{\left[ RNA \right]^{2}}{\left[ RNA \right]_{dimer}}=c^{⊖}e^{\frac{\Delta G}{RT}}$

${[RNA]}_{dimer}+\left[ RNA \right]\rightleftharpoons{[RNA]}_{trimer}\Longrightarrow K_{trimer}=\frac{[RNA]\left[ RNA \right]_{dimer}}{\left[ RNA \right]_{trimer}}=\frac{\left[ RNA \right]^{3}}{K_{dimer}\left[ RNA \right]_{trimer}}$

${[RNA]}_{trimer}+\left[ RNA \right]\rightleftharpoons{[RNA]}_{tetramer}\Longrightarrow K_{tetramer}=\frac{[RNA]\left[ RNA \right]_{trimer}}{\left[ RNA \right]_{tetramer}}=\frac{\left[ RNA \right]^{4}}{K_{dimer}K_{trimer}\left[ RNA \right]_{tetramer}}$

And so on to infinite *n*-mer. The total molar concentration of [RNA] would then be given by:

$${[RNA]}_{tot}=\left[ RNA \right]+2{[RNA]}_{dimer}+3{[RNA]}_{trimer}+4{[RNA]}_{tetramer}\ldots$$

Which can be rewritten as:

$${[RNA]}_{tot}=\left[ RNA \right]\left( 1+\frac{2[RNA]}{K_{dimer}}+\frac{3\left[ RNA \right]^{2}}{{K_{dimer}K}_{trimer}}+\frac{{4\left[ RNA \right]}^{3}}{{{K_{dimer}K}_{trimer}K}_{tetramer}}+\ldots\right)$$

Under the simplifying assumption that all dissociation constants are equal, so that growing an aggregate does not introduce any bonus nor penalty, we can define some useful constants:

$$x=\frac{[RNA]}{K}$$

$$L=\frac{{[RNA]}_{tot}}{K}$$

And rewrite $L$ as follows:

$$L=x\left( 1+2x+3x^{2}+4x^{3}+\ldots\right)$$

Since in such self-aggregating systems we find that *x* < 1, the series in parenthesis can be solved, obtaining the following:

$$L=\frac{x}{\left( 1-x \right)^{2}}$$

Which can be expanded as a quadratic equation in $x$, yielding the positive solution:

$$x=\frac{2L+1-\sqrt{4L+1}}{2L}$$

Finally, the fraction of monomers can be written as:

$$\frac{[RNA]}{{[RNA]}_{tot}}=\frac{x}{L}=\frac{2L+1-\sqrt{4L+1}}{2L^{2}}$$

By setting ${[RNA]}_{tot}=20 \mu M$ and $\Delta G=-7 kcal/mol$, the expected aggregated fraction ($\theta$) inside the cell can easily be computed as:

$$\theta=1-\frac{2L+1-\sqrt{4L+1}}{2L^{2}}=1-\frac{\frac{2\left[ RNA \right]_{tot}}{c^{⊖}e^{\frac{\Delta G}{RT}}}+1-\sqrt{\frac{4\left[ RNA \right]_{tot}}{c^{⊖}e^{\frac{\Delta G}{RT}}}+1}}{2\left( \frac{\left[ RNA \right]_{tot}}{c^{⊖}e^{\frac{\Delta G}{RT}}} \right)^{2}}\approx0.7$$

2. **Effect of different *k_on_* values on the simulation**

Our simulations have been run with a fixed *k_on_* equal to 10^7^ M^-1^s^-1^ based on a recent assessment on short RNA oligonucleotides (4). Literature values tend to vary depending on the specific experimental design and methodology used to measure these processes, leaving open the question of the potential effect of variation in this parameter on the simulations’ outcome.

In our simulation scheme, we have bound *k_on_* and *k_off_* to the equilibrium constant for the interaction as computed by ViennaRNA (2). This means that lowering the *k_on_* would lower the *k_off_* by the same amount, and vice-versa. The net effect that we would expect from such change would be the overall slow-down or speed-up of the systems’ dynamics, but not a change in their steady state properties. This was directly tested on a smaller system consisting in the 866 sequences that are part of the cluster shown in Fig. 3B. Our simulations revealed that the properties at steady state are not affected by *k_on_*, and changing the hybridization rate only delays the time needed to approach such state (Fig. S2).

3. Effect of different energy thresholds on the simulations

The prediction of interactions among *E. coli* mRNAs yielded ~28 million valid hybridization. Accounting for all of them in our simulations would have made them computationally untreatable. At the same time, most of these interactions are extremely weak and unlikely to participate in any interactions at any given time. To make the simulations feasible, we decided to apply an energy threshold, cutting off such weak interactions.

To determine the effect of thresholding on our simulation’s outcome, we tested two more stringent conditions, with a net observed effect of a reduction in the extent of aggregation (Fig. S3). To understand why this happens, we can look into the interactions data. In our system, we retrieved ~70,000 hybridizing stretches with energies equal or lower than -10 kcal/mol. For two molecules at 2.5 nM, this binding energy would correspond to a binding probability at equilibrium roughly larger than 3%. Although this probability may seem already small, it would be a mistake to disregard even weaker interactions as irrelevant: including stretches with hybridization energy as weak as -8 kcal/mol (binding probability ≳ 1%) increases the total number of potentially hybridizing stretches up to ~500,000, and including interactions as weak as -7 kcal/mol (binding probability ≳ 0.02 %) increases the number to ~1,300,000. Even if binding events between weaker and weaker stretches are unlikely, their abundance grows so dramatically that these small binding probabilities end up becoming relevant. This is immediately apparent when comparing the effect of these different energy thresholds on the simulations’ outcome as shown in Fig. S3, where pushing the boundary to weaker interactions increases both the systems connectivity and pairings.

Even though we used a -7 kcal/mol cutoff to get rid of extremely weak interactions, this analysis suggests that including them could further increase the connectivity of the system, driving it towards more interactions and cluster formation. In this sense, our simulation represents a conservative estimate of the degree of intermolecular hybridization in the system.

4. Sequence-shuffle: analysis pipeline

For each among the top 100 most abundant mRNAs, we generated 100 variants using either a codon-shuffle or dinucleotide-shuffle algorithm. The first approach has the advantage of keeping the identity of the encoded protein intact, while completely losing mRNA similarity. On the contrary, the second approach maintains the same nucleotide composition and total folding potential, while losing its encoded information.

For each sequence, we compared the value of the statistic of interest with the values of the same statistic in the pool of sequence-shuffled counterpart. To compare these values, we employed a custom non-parametric approach (Fig. S10). Briefly, as a null-hypothesis we expected the natural sequences to have features comparable to the randomized ones. This could be quantified by computing the one-tail probability (P-value) of generating a random sequence that has a lower statistic than the natural sequence. In the presented case study, we counted the fraction of randomized sequences that had a lower (more negative) MFE than the naturally occurring mRNA.

Once each gene was tested, the P-values were pooled together, and a probability distribution was generated. The shape of this distribution can be one out of three: (1) if the P-values tend to be small, the probability distribution will be skewed to the left; (2) if the P-values are not significantly larger or smaller than 0.5, the distribution will be uniform, and finally (3) if the P-values tend to be large, the probability distribution will be skewed to the right. To determine what scenario better described out statistics, the skewness of the probability distribution was quantified. This was achieved by computing the integral (∑, total area under the curve) of the cumulative probability distribution of the P-values. This parameter ∑ can vary between 0.0 and 1.0, with ∑ close to zero signifying that the statistics is shifted to large values, ∑ close to one signifying that the statistic is shifted to small values, and finally ∑ close to 0.5 signifying the lack of a preferential shift.

Although our method neatly quantifies the differential behavior of natural sequences and the existence of selective pressure, the entity of such difference is not intuitively caught by the ∑ metric. To provide a more intuitive measure for the magnitude of the difference between the naturally occurring sequences and their randomized counterparts, we computed the *equivalent shift for a normal distribution*. In other words, we determine the magnitude of the shift that would have been observed if both null-statistic and natural statistic had followed normal distributions. Briefly, let’s consider a null-statistic that follows a standard distribution, and assume that a statistic of interest follows a normal distribution as well, only shifted by some amount (𝛿) and potentially stretched (thus having different variance). We can sample multiple values for the statistic and analyze them with our pipeline to compute their P-values against their reference null-distribution (standard distribution). This process can be repeat by modifying 𝛿 and the distribution variance until they yield a probability distribution for P-values (and thus ∑) that matches the observed one. These 𝛿 are reported as converted values for ∑.

5. Sequence-shuffle: results

For the MFE, we found that natural sequences in *E. coli* fall in the first scenario, with a large ∑ equal to 0.9 (Fig. 5A) meaning that naturally occurring mRNAs can typically achieve a more stable folded state than their randomized counterpart. It could be argued that such result may be the lucky outcome of a random sampling process in the large protein sequence space, and not truly the effect of natural selection at play. To ensure this was not the case, we performed a bootstrap analysis by sampling 1000 random reduced genomes, each including the same top 100 genes encoding the same proteins with one among the 100 possible CDS variants previously generated. With this approach, we found that not even one of the 1000 random genomes (*p < 10^-3^*) produces a ∑ as extreme as the natural reduced genome.

Although our method neatly quantifies the differential behavior of natural sequences and the existence of selective pressure, the entity of such difference is not intuitively caught by the ∑ metric. To improve its readability, this value was converted to an equivalent shift in standard deviations (𝛿) of the random distribution (see Supplementary Note 4), finding that the MFE for natural sequences were shifted by a staggering -2.0 𝛿.

The MFE is a useful metric related to the potential of aggregation for mRNAs, but it does not provide a complete and satisfying description. To better characterize this potential, we proceeded with the analysis of two more parameters. First, we analyzed the length of the longest Unpaired Segment (US) that could be exposed upon folding for a given CDS in the reduced genome (Fig. S11(b)). Similarly to the MFE, we found that this statistic is significantly shifted to low values (*p < 10^-3^*), yielding an ∑ = 0.71 (equivalent to -0.9 𝛿), signifying that natural sequences have a tendency to expose shorter single stranded regions, reducing the chance for extensive complementary and spurious intermolecular interactions.

Finally, we directly analyzed the hybridization energy (previously defined as ${\Delta G}_{min}$) for the most stable duplex that can be formed among any out of the 5050 unique pairs of the top 100 most abundant mRNAs in *E. coli* (Fig. 5C). This metric more directly measures the potential for aggregation, and it revealed once again a significant (*p < 10^-3^*) differential behavior for natural sequences, yielding an ∑ = 0.36 (equivalent to +0.6 𝛿). Importantly, a ∑ lower than 0.5 means that these energies are not shifted to lower (more negative values) as observed for the MFE, but the exact opposite; natural sequences tend to have larger (more positive) hybridization energies, and thus a weaker potential for intermolecular interactions.

These results depict the genome of *E. coli* as subject to selection pressure to produce its proteins utilizing mRNA sequences optimized to avoid cross-interactions. One caveat of this approach is that by swapping codons and introducing synonymous mutations, we also inevitably altered the CG content of the RNA sequences, which can change base-pairing potential and influence the outcome of our analysis.

As an alternative randomization method that does not suffer from this issue, we employed dinucleotide-preserving shuffles. Sequences generated this way keep length and dinucleotide composition constant while losing sequence-encoded information, so that the shuffled mRNAs will not encode for the same protein anymore. The strength of this approach consists in conserving the base-pairing potential by preserving nearest-neighbor stacks (7) – something that would not be true for simple nucleotide-preserving shuffles –, while randomizing higher-order sequence features.

For this alternative analysis (Fig. 5A-F), we once again worked on the reduced genome containing the top 100 most abundant mRNAs, and shuffled their sequences using the dinucleotide shuffling algorithm by Altschul-Erikson (8). This study confirmed our previous observations, so that both MFE (∑ = 0.84, equivalent to -1.4 𝛿), length of the longest US (∑ = 0.68, equivalent to -0.8 𝛿) and ${\Delta G}_{min}$ (∑ = 0.39, equivalent to -0.4 𝛿) are significantly (*p < 10^-3^* for all cases) different from randomized genomes having the same dinucleotide composition and base-pairing potential.

6. Quantifying ViennaRNA accuracy

In this work we have relied on ViennaRNA (2) to predict the folded structure of *E. coli* mRNAs. To determine whether these predictions could be used as proxies for *in vivo* mRNA accessibility, we compared computed values with experimental DMS-Seq data (9) retrieved from the RASP atlas(10) for a set of randomly picked *E. coli* sequences. Each entry allows to infer RNA accessibility by providing reactivities of cytosines and adenines to the alkylating agent DMS at the single nucleotide resolution.

DMS experiments generate a list of reactivities to the alkylating agent that can be normalized over a continuous interval from zero to one but unfortunately do not provide a binary output for paired/unpaired nucleotides. For any DMS experiment, a reactivity threshold has to be picked to determine what nucleotides can be considered DMS-paired (reactivity below the threshold) or DMS-unpaired (reactivity above the threshold).

To evaluate whether ViennaRNA agreed with experimental evaluation of mRNA structures *in vivo*, we computed the expected base-paired probability of our transcripts of choice and then split their nucleotides into “weakly paired” and “strongly paired” based on their *P_i_* values as previously defined.

For each subset of “weakly paired” and “strongly paired” nucleotides as predicted with ViennaRNA, we computed the fraction of bases whose state agrees with DMS assessment as we varied the DMS-reactivity threshold. This approach yields “True Paired Rate” vs “True Unpaired Rate” monotonously decreasing functions, highlighting the compromise between high sensitivity and specificity. The integral of the area for these curves (AUC) directly informs on the agreement between the two methods. In all cases analyzed here, we found that the AUC was larger than 0.5, with many cases showing exceptionally high concordance (Fig. S11).

7. Simulation scheme: additional details

The specific dehybridization rates (*k_off_*) for each pair of unpaired segments was computed from *k_on_* and ${\Delta G}_{min}$ using Equation (3). By binding *k_on_* and *k_off_* through the equilibrium constant, we made sure that the properties at steady state - the main focus of our downstream analysis - are independent of our specific choice of rates (see Supplementary Note 2).

Reported hybridization rates (*k_on_*) for nucleic acids in literature span between 10^6^ to 10^7^ M^-1^s^-1^(58, 59, 18). Although length, CG content (18), coaxial stacking (60) and defects (mismatches, bulges) (61) influence the hybridization rate, their effect on *k_on_* is typically modest when compared to their impact on the dehybridization rate (*k_off_*). We therefore fixed *k_on_* = 10^7^ M^-1^s^-1^ in line with the nucleation-limited values reported for short RNA oligonucleotides (18). This choice of parameters yielded dissociation timescales that could be as long as ~2 years for the most stable duplexes in our set.

Because the Gillespie algorithm tracks individual molecules, we converted the concentration-dependent hybridization rate *k_on_* to an effective per-pair reaction rate. Given the *E. coli* intracellular volume of 6.7 x 10^-10^ µL (21), the effective concentration of a single molecule is approximately 2.5 nM, yielding a constant hybridization rate of 2.5 x 10^-2^ s^-1^ for any pair of available unstructured stretches.

Noticeably, microscopic hybridization rates assume no competition. Since the median unstructured stretch can interact with ~70 partners, the apparent *k_on_* that can be measured experimentally for one chosen target is going to be much smaller than the theoretical value. This is because any trajectory that would lead to hybridization against a specific target will most likely pass through several events of hybridization and dehybridization with off targets. This behavior is conveniently captured for free by the stochastic simulation, and it is an emergent properties that does not require any additional inputs or modifications (18).

The simulation proceeds one event at a time, consisting of either the hybridization of two unstructured stretches on two mRNAs to form a new duplex, or the dehybridization and dissolution of an existing one. Since low hybridization energies correspond to fast dynamics, weak duplexes form and dissolve at high frequency. Including these events would force the simulation to spend the vast majority of its time processing these transient interactions. Simulating all possible pairwise interactions would be computationally prohibitive, and it is made possible thanks to the thermodynamic cut-off previously introduced. This practical compromise ensured computational feasibility while retaining the most stable and likely consequential interactions. Importantly, a higher threshold will not only be more realistic by accounting for more potential events, but it will likely lead to even more interaction (see Fig. S3 for the evaluation of different thresholds).

**Supporting Information**

- Figures S1 to S12
- Tables S1 to S2
- SI References

| Table of Content | Page |
| --- | --- |
| Fig. S1, Pipeline for predicting mRNA intermolecular interactions | 11 |
| Fig. S2, Effect of different *k_on_* on the simulations’ outcome | 12 |
| Fig. S3, Effect of different energy thresholds on the simulations’ outcome | 13 |
| Fig. S4, Clusters disassembly dynamic | 14 |
| Fig. S5, Study of clusters topology | 15 |
| Fig. S6, *E. coli* mRNA readily aggregates *in vitro* | 16 |
| Fig. S7, Experimental study of polynucleotide aggregation | 17 |
| Fig. S8, Length as predictor of mRNA aggregation | 18 |
| Fig. S9, Genes overlap between sequencing and simulation | 19 |
| Fig. S10, Statistical analysis of shuffled mRNA sequences in *E. coli* | 20 |
| Fig. S11, Folding of *E. coli* mRNAs by ViennaRNA | 21 |
| Fig. S12, Simulation and sequencing abundance comparison | 22 |
| Table S1. Minimum Folding Energy for native abundant mRNAs in *E. coli* compared to matching shuffled sequences. | 23 |
| Table S2. Longest unpaired stretch for native abundant mRNAs in *E. coli* compared to matching shuffled sequences. | 25 |
| SI References | 27 |

Fig. S1, Pipeline for predicting mRNA intermolecular interactions

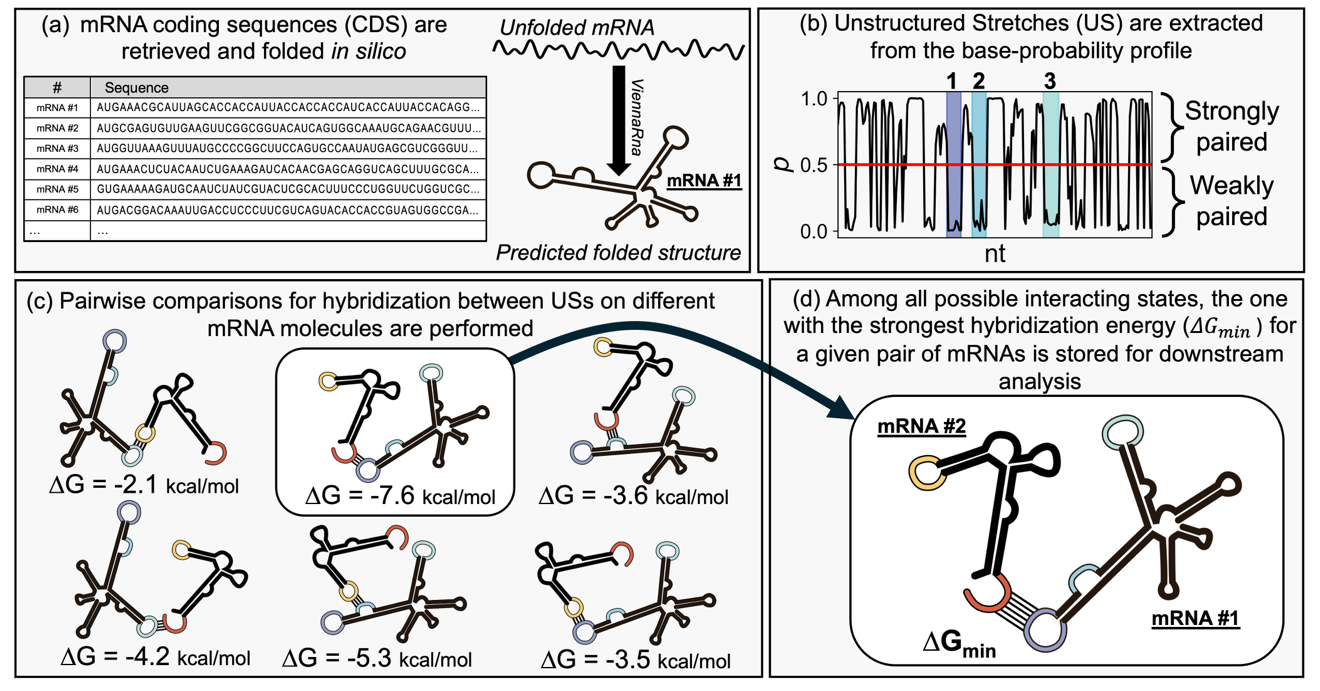

(a) All annotated mRNA sequences are retrieved from a transcriptomic compendium (1) and folded using ViennaRNA (2). (b) The regions corresponding to long stretches of nucleotides (≥ 7 nt) having low base-pairing probability (*p* < 0.5) are annotated as Unstructured Stretches and highlighted as shaded area in the graph. In the example for mRNA #1, three USs satisfy our criteria. (c) For all pairs of mRNAs in the dataset, the free energy for hybridization for all their USs is calculated. In the example, mRNA #1, having three USs, is compared with mRNA #2, having 2 USs. (d) The most stable interaction per pair of mRNAs is stored for the simulation. In the example, the most stable interaction leads to a free energy change of -7.6 kcal/mol which is thus the ${\Delta G}_{min}$ of this specific pair of sequences.

**Fig. S2,** **Effect of different *k_on_* on the simulations’ outcome**

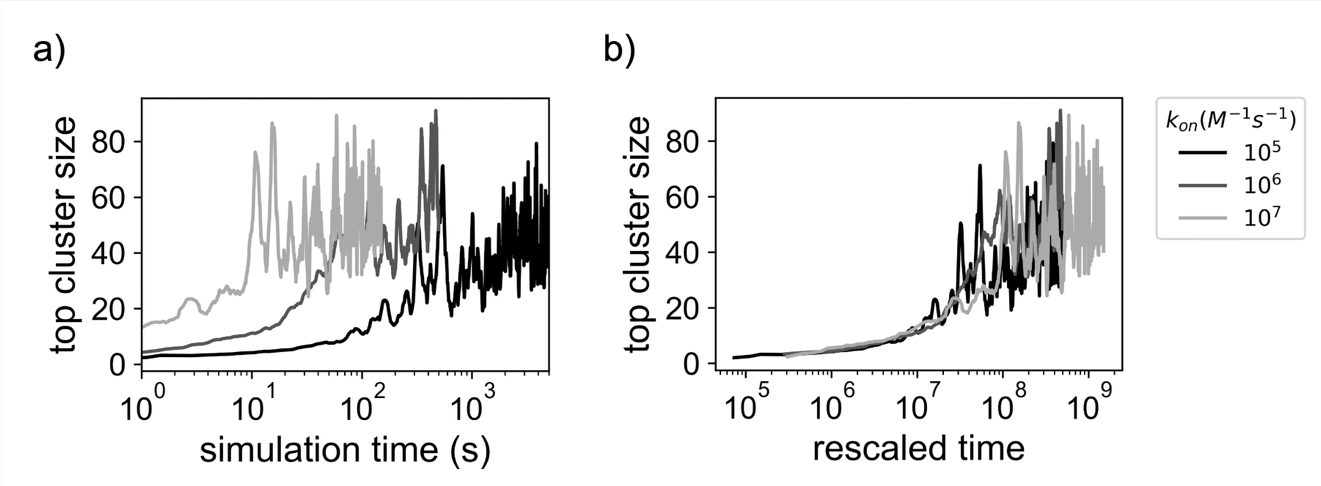

Results on simulation performed on a small system consisting in the 866 sequences that are part of the cluster shown in Figure 3B. Varying the chosen hybridization rate parameter, we found that a) lower *k_on_* values delay the approach to a comparable steady state. b) Normalizing the time axis by *k_on_* leads to overlap between the different trajectories, confirming that changes in rates have the overall effect of proportionally stretching or shrinking the time scale.

**Fig. S3,** **Effect of different energy thresholds on the simulations’ outcome**

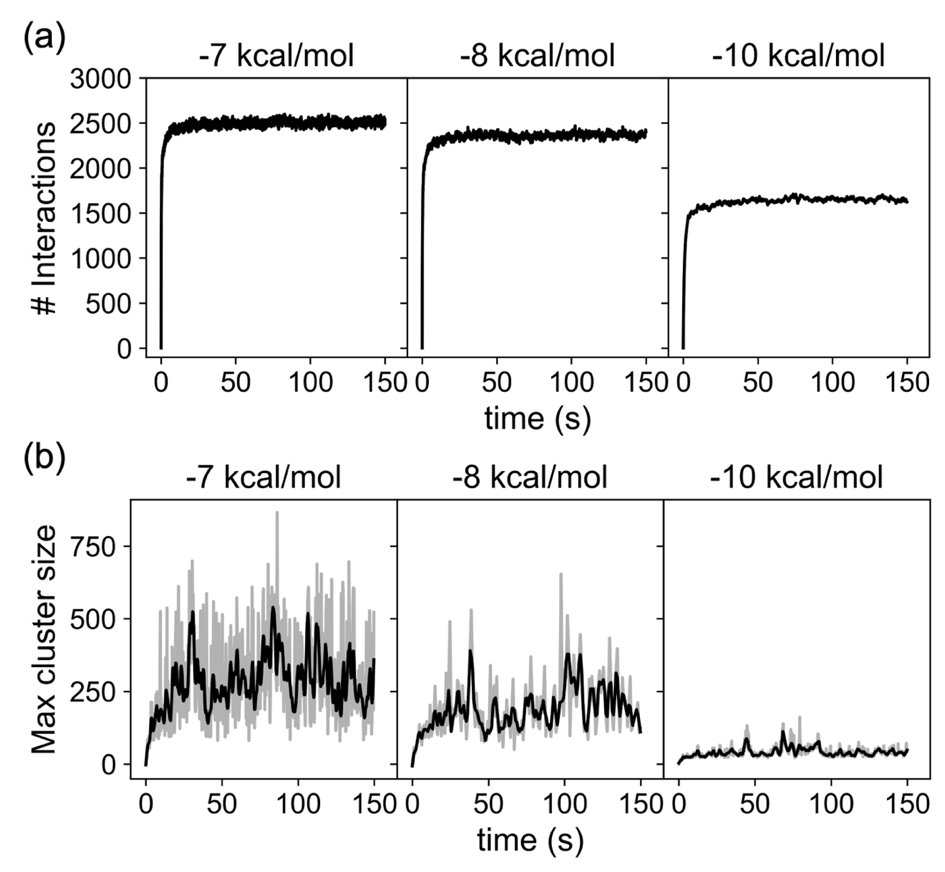

Alternative simulations were performed by progressively keeping only the most stable (and scarce) interactions. The (a) total number of interactions and (b) largest cluster size were found to decrease if only the most stable interactions are allowed in the simulation.

Fig. S4, Clusters disassembly dynamic

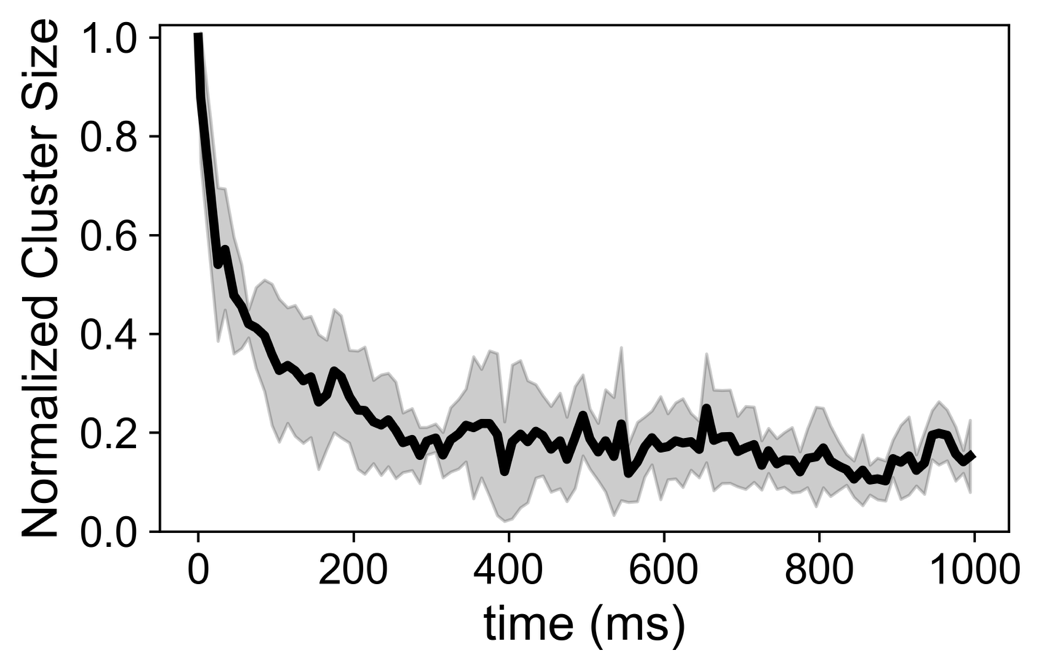

Interactions between RNA in the simulation are short-lived, with large clusters dissolving in less than a second. The trace shows the normalized mean trajectory of five clusters with initial size ≳ 700. All traces were aligned at time zero corresponding to the moment their size reached a local maximum.

**Fig. S5, Study of clusters topology**

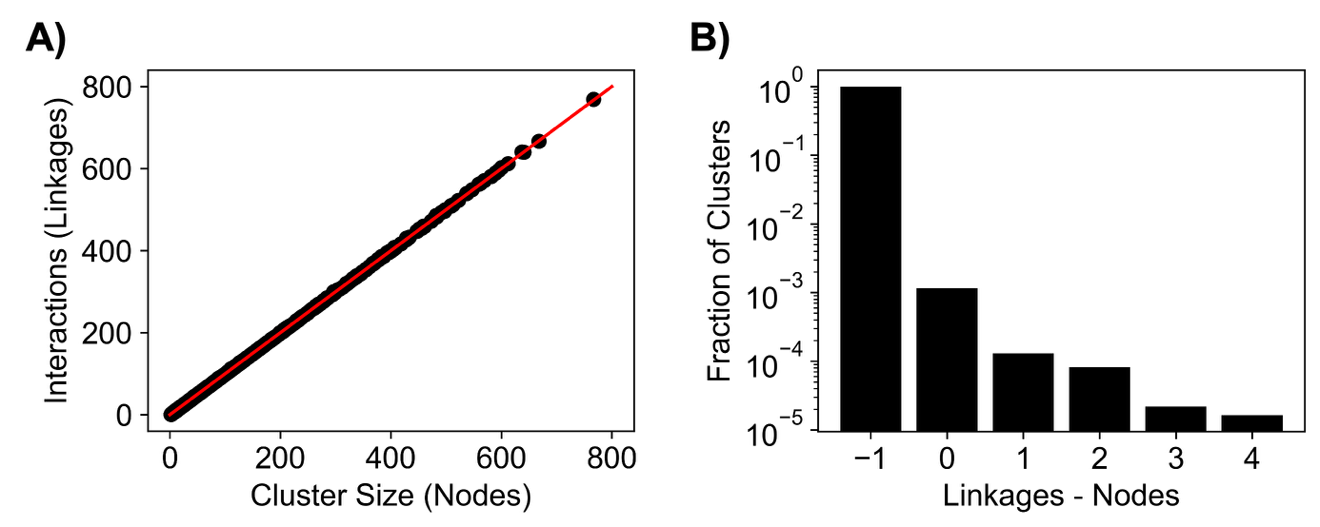

A) To determine the topology of the clusters formed by RNA molecules in our simulations, we systematically computed their size and number of interactions at steady state finding that these numbers are comparable. This matches the expectations from a tree-like network without loops, where the number of interactions (linkages) is equal to the number of elements in the cluster (nodes) minus 1. Instead, in highly crosslinked structures, we would have expected the difference between number of linkages to be larger than the number of nodes.

B) By computing the likelihood of observing clusters with various degrees of crosslinking in our simulations, we found that more than 99% are tree-like, and a small fraction of clusters has a modest degree of crosslinking.

Fig. S6, *E. coli* mRNA readily aggregates *in vitro*

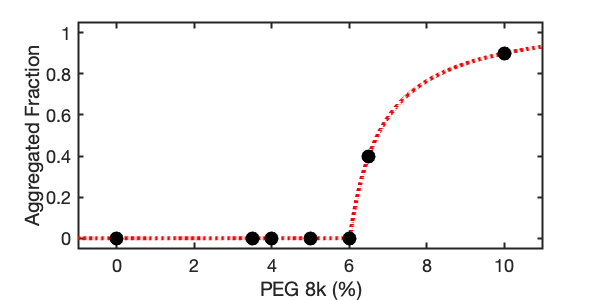

The addition of PEG causes *E. coli* mRNA to readily aggregate *in vitro*. The study has been performed using 5 ng/μL RNA, 1 M NaCl and SYBR Gold (1x). Quantification of RNA in the aggregated fraction was determined using a Nanodrop 8000 (Thermo Scientific). Dotted line is a guide for eye.

Fig. S7, Experimental study of polynucleotide aggregation

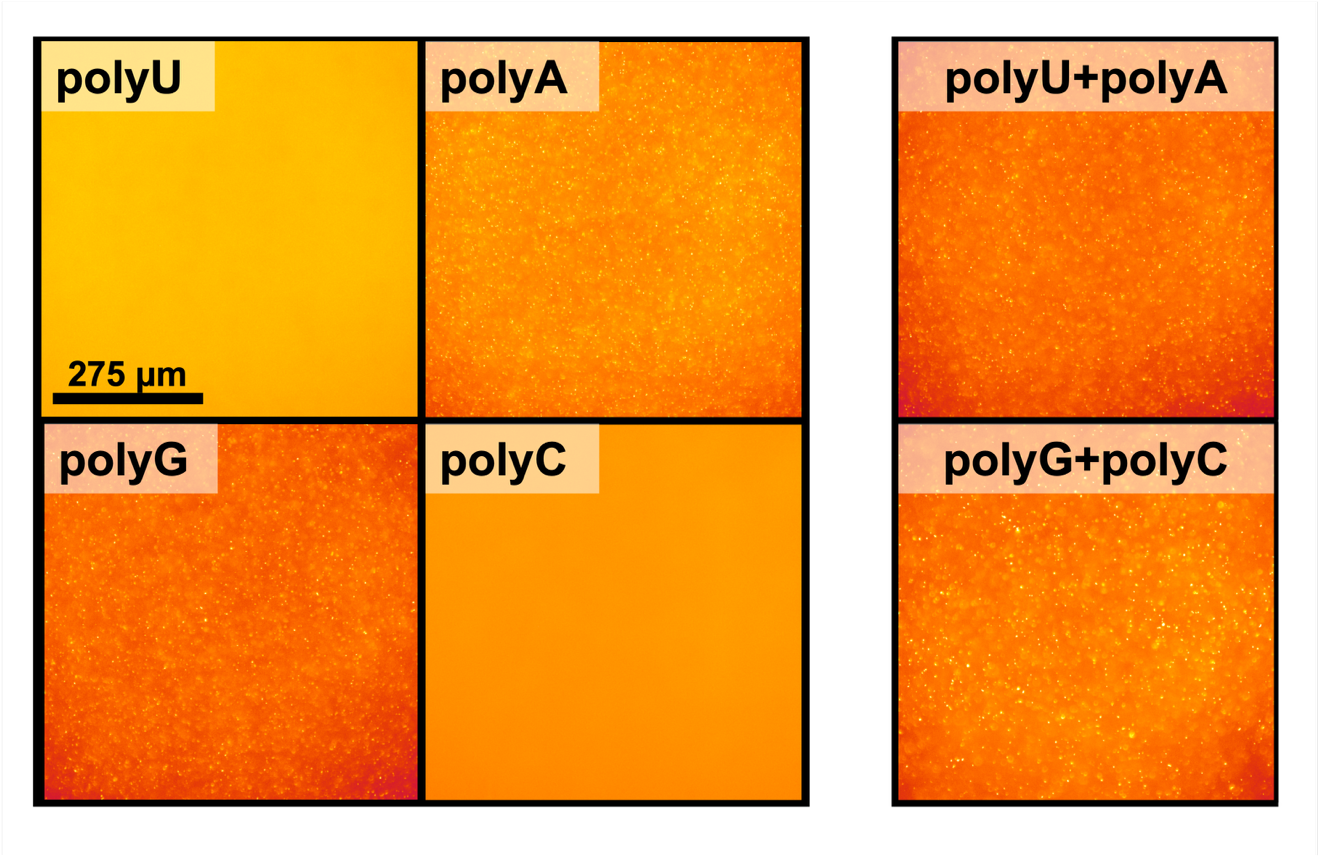

Homopolymeric RNA aggregation *in vitro*. Fluorescent micrographs for samples containing individual homopolymeric RNAs or pairs of complementary ones. Our computational model predicts widespread aggregation mediated by base pairing between exposed unstructured stretches of folded mRNAs. To evaluate if attractive intermolecular interactions are also required for aggregation *in vitro*, we studied the behavior of homopolymeric RNAs (5 ng/μL) in 10% PEG, 1 M NaCl and 5x SyberGold.

We found that polyU and polyC did not form detectable aggregates, whereas polyA and polyG readily self-assembled, consistent with their known propensity for self-association (5, 6). Moreover, mixtures of complementary polynucleotides did formed aggregates, showing that crowding alone is insufficient to drive aggregation.

Fig. S8, Length as predictor of mRNA aggregation

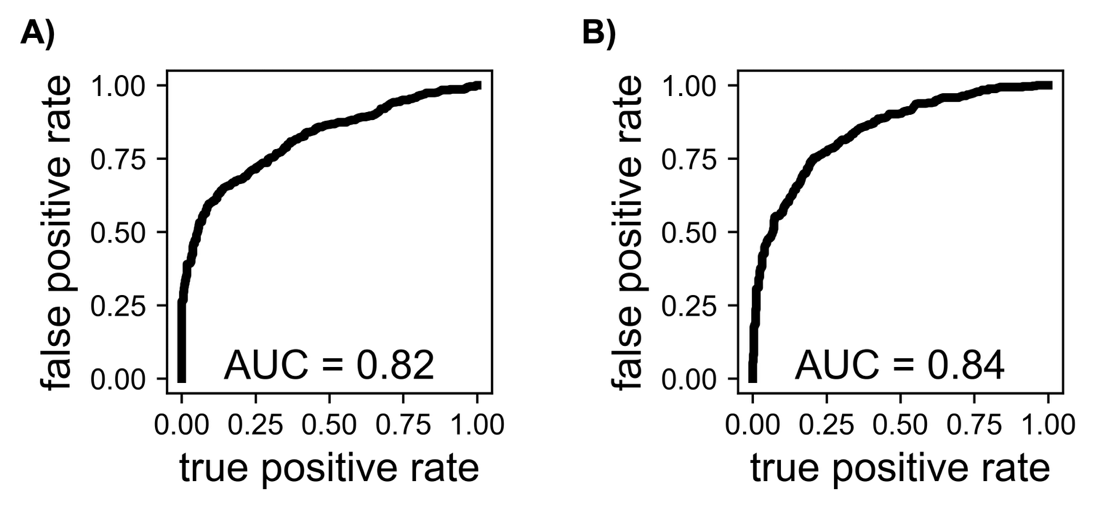

Aggregated E. coli mRNA, both *in vitro* and *in silico*, appeared enriched in longer transcripts. To quantify this effect, we calculated the fold change in normalized abundance for each transcript in the aggregated fraction relative to the input. Inspection of the top and bottom 500 transcripts ranked by fold change revealed a marked difference in their length distributions, suggesting that transcript length alone might predict enrichment in aggregates.

To formally test this hypothesis, we evaluated transcript length as a binary classifier distinguishing aggregated from non-aggregated transcripts. We constructed Receiver Operating Characteristic (ROC) curves by varying the length threshold and plotting the true positive rate against the false positive rate. The area under the ROC curve (AUC) quantifies classification performance, with 0.5 corresponding to random prediction and 1.0 to perfect discrimination.

In both experimental (A) and simulation (B) datasets, transcript length achieved an AUC greater than 0.8, indicating that length alone is a strong predictor of aggregation propensity.

Fig. S9, Genes overlap between sequencing and simulation

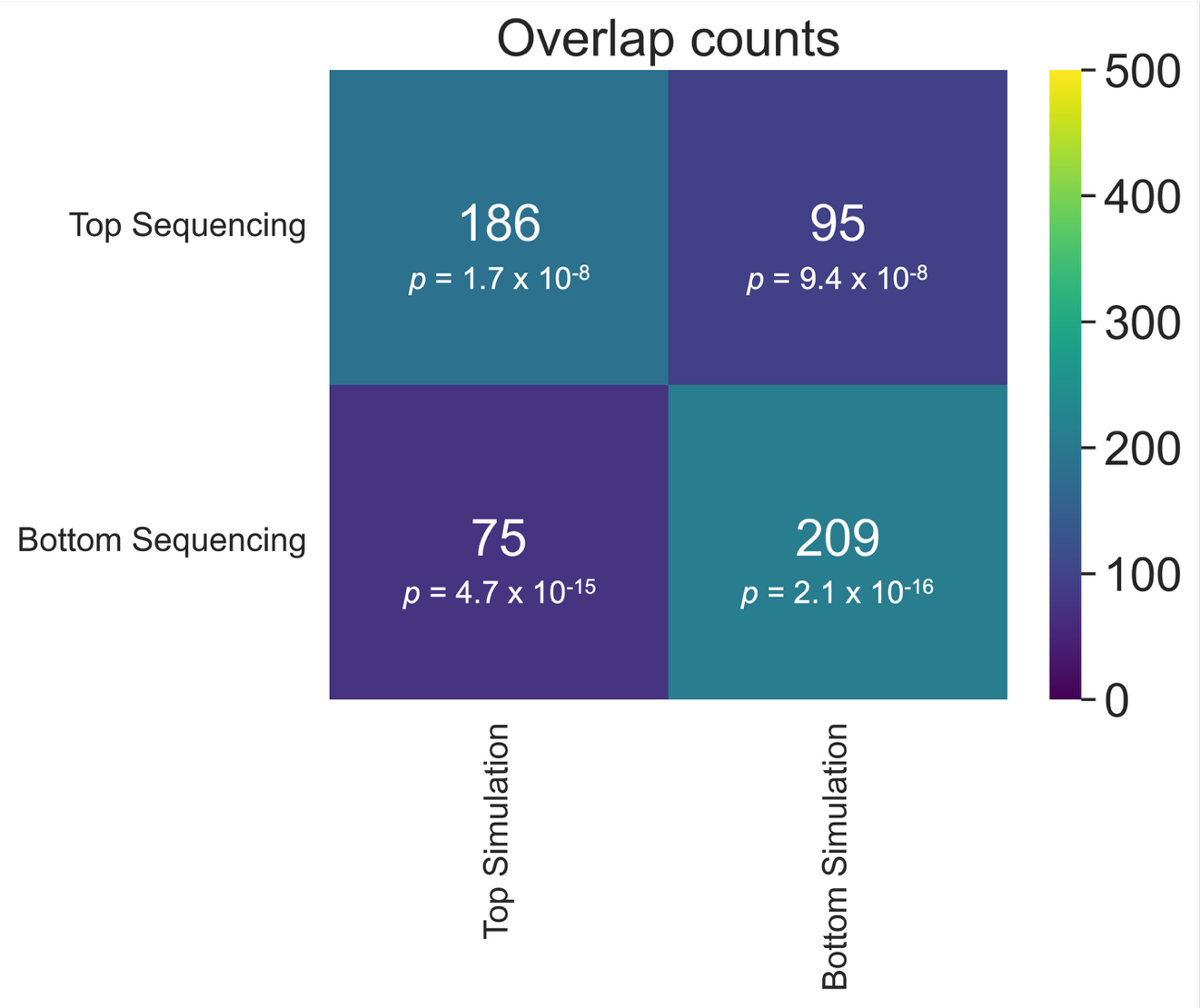

Our enrichment analysis based on simulation and sequencing data both showed that the top and bottom 500 transcripts (ranked by fold change) were respectively preferentially enriched and depleted for long sequences. To further corroborate the hypothesis of a shared underlying mechanism driving aggregation, we quantified the overlap between transcripts enriched (top 500) or depleted (bottom 500) in simulation and experiment. Under a null model in which transcripts are randomly assigned to these sets, a hypergeometric test predicts an average overlap of 138 RNAs (95% confidence interval: 121–154). In contrast, the enriched sets shared 186 transcripts, and the depleted sets shared 209 transcripts—substantially exceeding random expectation.

**Fig. S10, Statistical analysis of shuffled mRNA sequences in *E. coli***

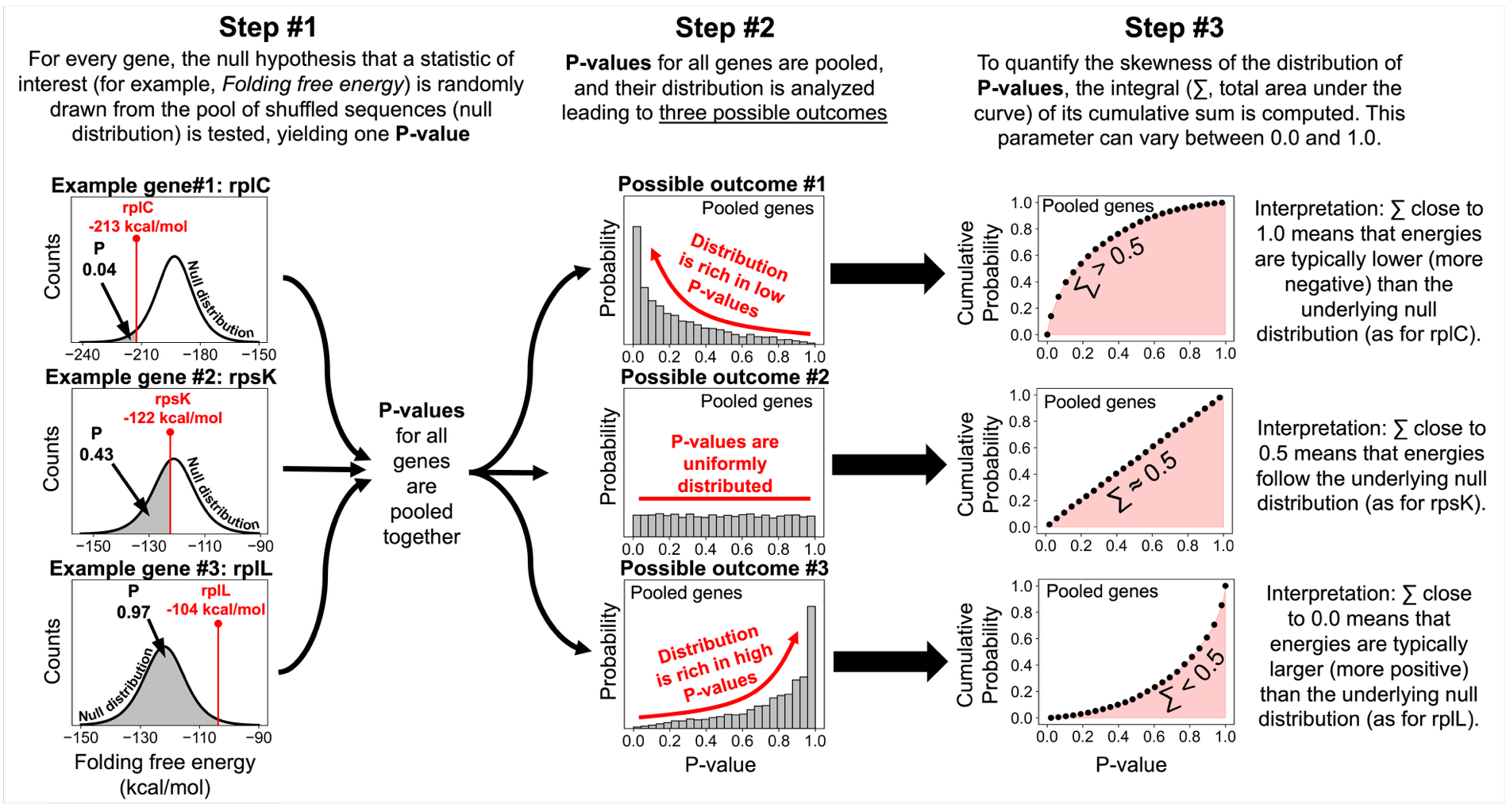

The analysis proceeds through three steps. **Step #1:** the null hypothesis that a statistic of interest (for example, minimum folding free energy - MFE) smaller than the natural one is randomly drawn from the pool of shuffled sequences (null distribution) is tested, yielding one P-value. **Step #2:** P-values for all genes are pooled and analyzed. Their probability distribution can assume one out of three possible forms, leading to three different outcomes. **Step #3:** To quantify the skewness of the distribution of P-values, the integral (∑, total area under the curve) of its cumulative sum is computed. This parameter can vary between 0.0 and 1.0, with discrepancies from 0.5 revealing sequence selection for the statistics being either smaller (∑ > 0.5) or larger (∑ < 0.5) than what expected just by chance.

**Fig. S11, Folding of *E. coli* mRNAs by ViennaRNA**

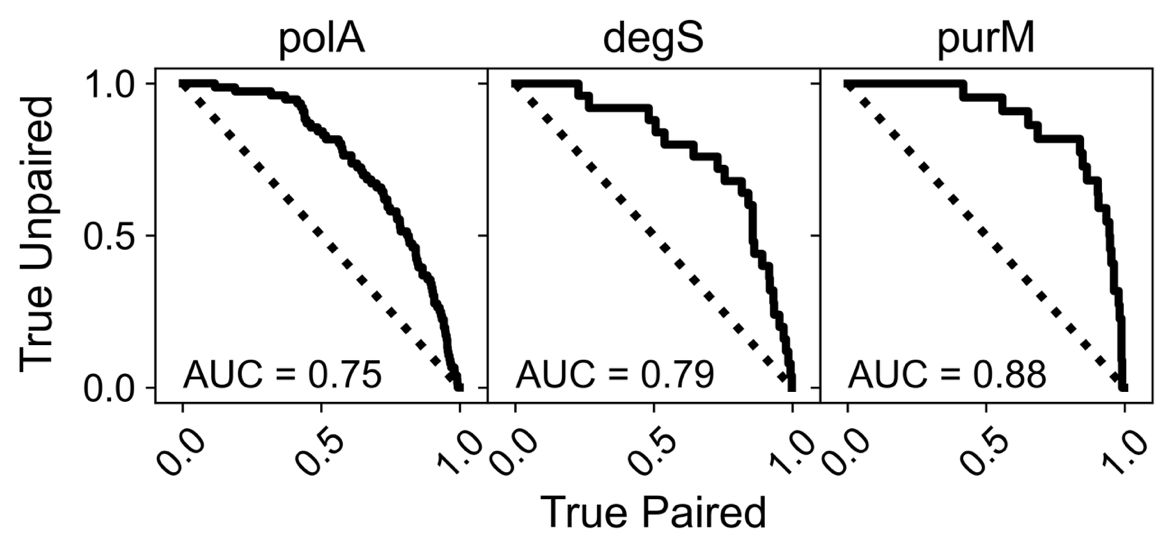

Folding of *E. coli* mRNAs by ViennaRNA is in good agreement with experimental assessment of RNA accessibility by DMS-Seq. (a, b, c) Examples of sequences yielding high agreement between the two methods.

Fig. S12, Simulation and sequencing abundance comparison

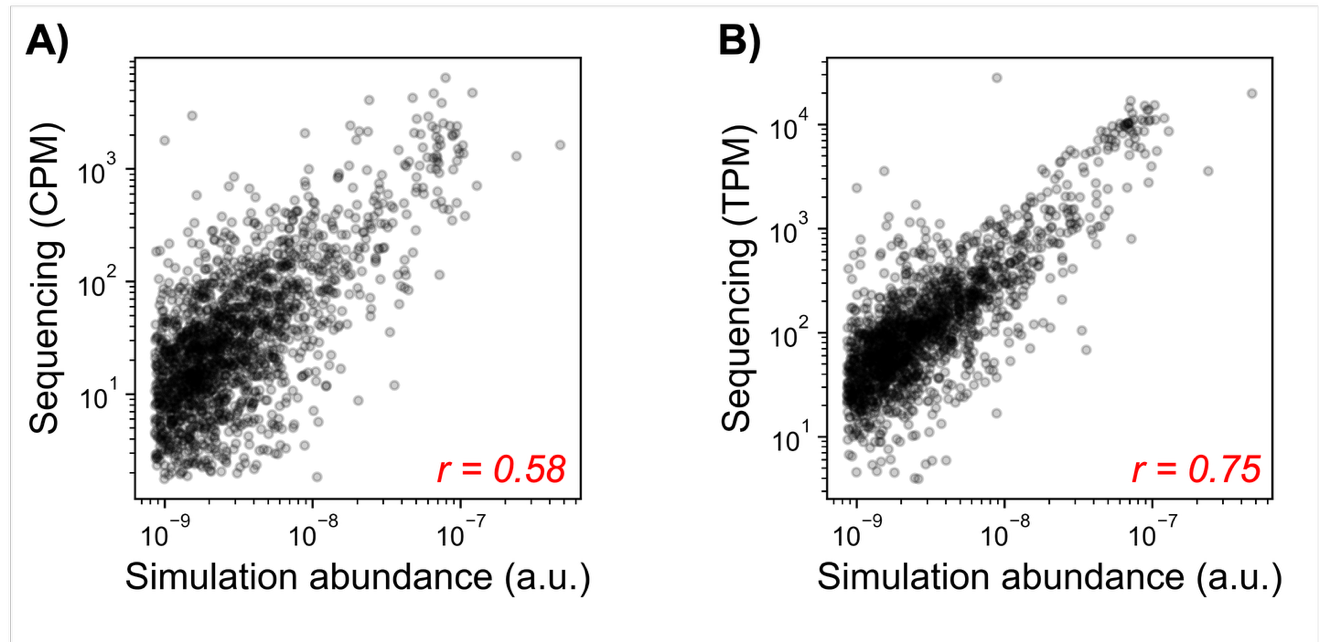

Sequences abundance from our sequencing experiments reported as either CPM (A) or TPM (B) versus the abundances used for our simulations (1). Pearson’s correlation coefficients are reported in red.

**Table S1.** **Minimum Folding Energy for native abundant mRNAs in *E. coli* compared to matching shuffled sequences.**

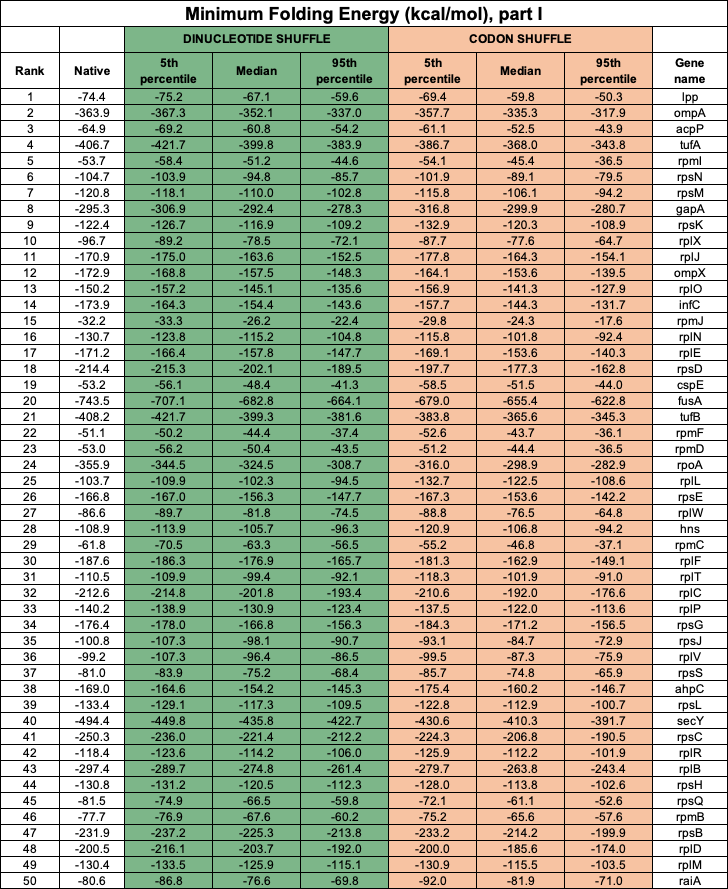

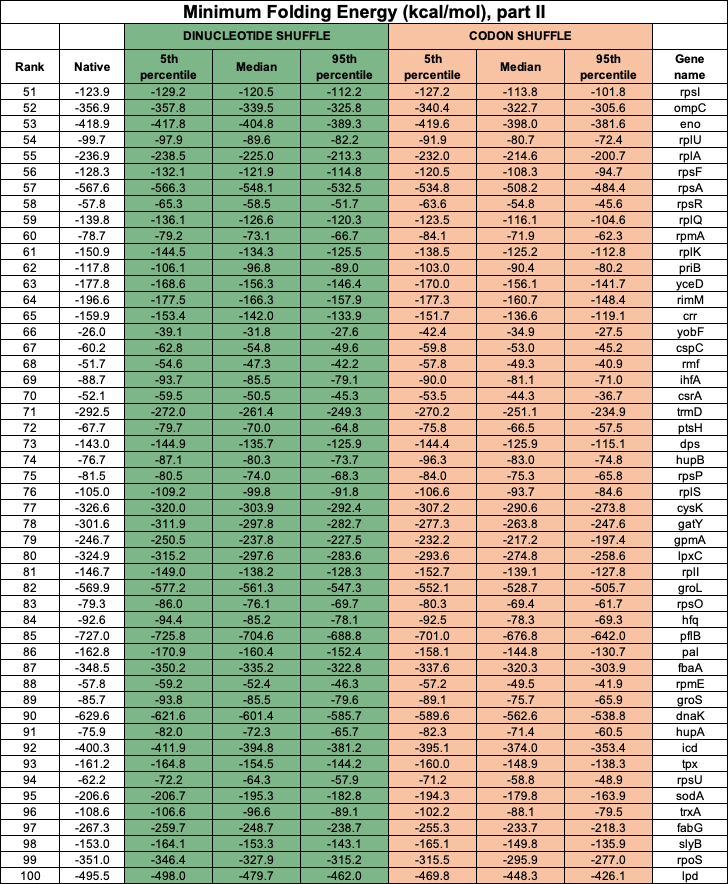

**Table S2. Longest unpaired stretch for native abundant mRNAs in *E. coli* compared to matching shuffled sequences.**

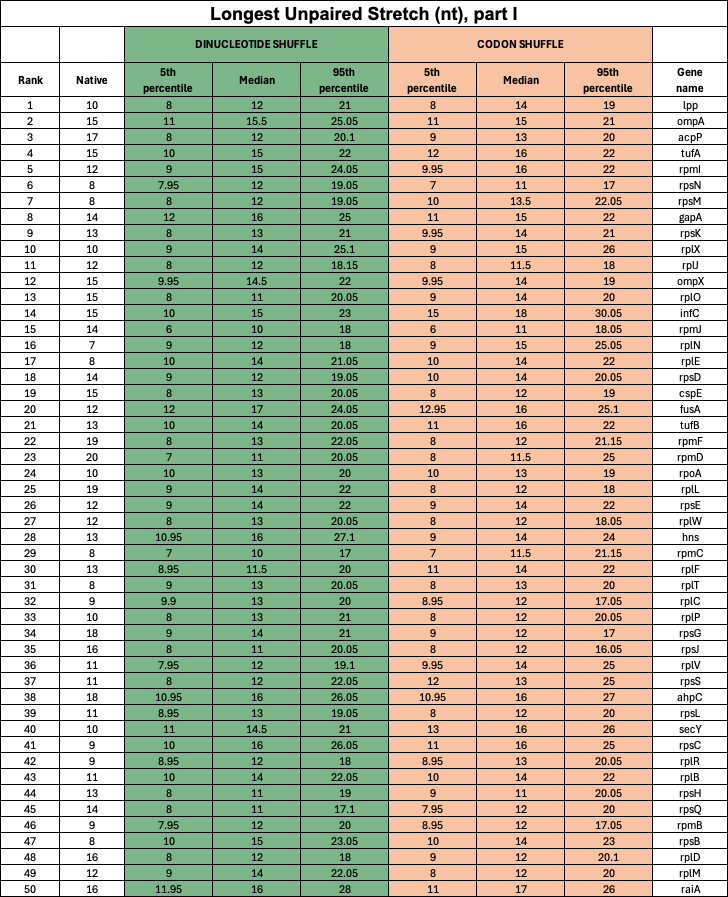

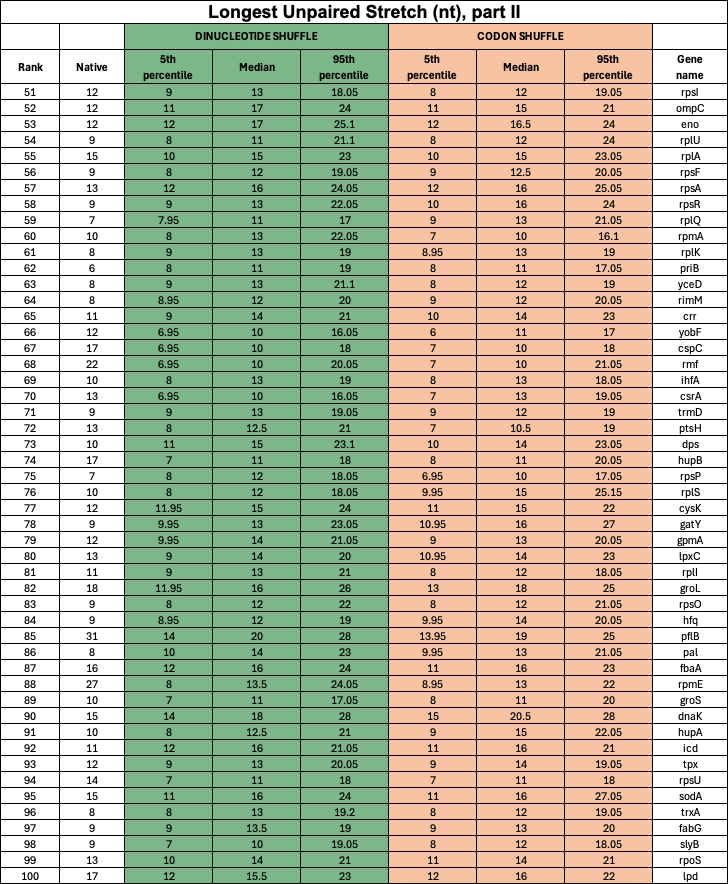
